# Visual and auditory brain areas share a representational structure that supports emotion perception

**DOI:** 10.1101/254961

**Authors:** Beau Sievers, Carolyn Parkinson, Peter J. Kohler, James M. Hughes, Sergey V. Fogelson, Thalia Wheatley

## Abstract

Emotionally expressive music and dance occur together across the world. This may be because features shared across the senses are represented the same way even in different sensory brain areas, putting music and movement in directly comparable terms. These shared representations may arise from a general need to identify environmentally relevant combinations of sensory features, particularly those that communicate emotion. To test the hypothesis that visual and auditory brain areas share a representational structure, we created music and animation stimuli with crossmodally matched features expressing a range of emotions. Participants confirmed that each emotion corresponded to a set of features shared across music and movement. A subset of participants viewed both music and animation during brain scanning, revealing that representations in auditory and visual brain areas were similar to one another. This shared representation captured not only simple stimulus features, but also combinations of features associated with emotion judgments. The posterior superior temporal cortex represented both music and movement using this same structure, suggesting supramodal abstraction of sensory content. Further exploratory analysis revealed that early visual cortex used this shared representational structure even when stimuli were presented auditorily. We propose that crossmodally shared representations support mutually reinforcing dynamics across auditory and visual brain areas, facilitating crossmodal comparison. These shared representations may help explain why emotions are so readily perceived and why some dynamic emotional expressions can generalize across cultural contexts.

## Introduction

Wherever there is music, there is movement (Kaeppler, 1978; Mehr et al., 2019; Savage, Brown, Sakai, & Currie, 2015). Not only are music and dance pervasive across the anthropological and ethnographic record, some languages use a single word for both (Baily, 1985; Trehub, Becker, Morley, & Trehub, 2015). The link between music and movement is present from early in development, with infants as young as 7 months using movement to resolve ambiguities in musical rhythm (Phillips-silver & Trainor, 2005). Further, communication of emotion through music and movement occurs across a range of dissimilar cultures (Fritz et al., 2009; Sievers, Polansky, Casey, & Wheatley, 2013; Trehub et al., 2015), although there are also many important cross-cultural differences in emotion expression, perception, and conceptualization (Gendron, Roberson, Vyver, & Barrett, 2014; Jack, Caldara, & Schyns, 2012; Jack, Sun, Delis, Garrod, & Schyns, 2016; Jackson et al., 2019; Margulis, Wong, Simchy-Gross, & McAuley, 2019; Yuki, Maddux, & Masuda, 2007). Here, we suggest that the link between music and movement may result from fundamental similarities in how music and movement are structured, perceived, and represented in the brain.

Supporting this account, preliminary research suggests that emotional music and movement can share structural features across cultures. In both the United States and a small-scale society in rural Cambodia, angry music and movement are both fast and move downward, peaceful music and movement are both slow and move upward, and so on (Sievers, Lee, Haslett, Wheatley, & Wheatley, 2019; Sievers et al., 2013). Though suggestive, shared features do not fully explain the pervasive, experiential link between music and movement. Here, we examine a possible explanation: Different sensory areas of the brain may share a representational geometry (Kriegeskorte & Kievit, 2013), such that differences between sensory features and perceived emotions are represented by matched differences in patterns of neural activity, putting music and movement in comparable, task-relevant terms.

We tested two related main hypotheses concerning both *where* and *how* music and movement are represented in the brain. (H1) The *separate regions, shared representations* hypothesis: that separate, modality-specific, auditory and visual areas use a shared representational geometry. (H2) The *supramodal region* hypothesis: that a supramodal area (or areas) uses a single representational geometry for both auditory and visual stimuli. Note that (H1) does not require patterns of activity in auditory and visual brain regions to be identical in every respect, as each sensory region likely represents modality-specific features. Evidence that the representation of music in auditory regions is very similar to the representation of movement in visual regions would support the separate regions, shared representations hypothesis (H1). By contrast, evidence of a single region that represents both music and movement using the same representational geometry would support the supramodal region hypothesis (H2). Importantly, (H1) and (H2) are not mutually exclusive, and while previous research has provided support for (H2) (2010), the status of (H1) remains unknown.

Further, we asked how representations of perceived emotion in music and movement were organized, testing two auxiliary hypotheses. (A1) The *simple features* hypothesis: that sensory brain regions represent emotional stimuli in terms of differences in simple stimulus features, without respect to how those features may later be inferentially processed to yield emotion judgments (by e.g., simulation theory, Gordon, 1986; or theory theory, Gopnik & Wellman, 1994). (A2) The *environmental conjunctions* hypothesis: that sensory representations of emotional stimuli closely track emotion judgments, suggesting that the human perceptual system may directly represent latent configurations of stimulus features associated with emotion content. These task-relevant representations may act as a shortcut, reducing the need for downstream inferential processing (Gallagher, 2008). Evidence that sensory representations fit a model based on stimulus features would support the simple features hypothesis (A1), while evidence that sensory representations fit a model based on emotion judgments would support the environmental conjunctions hypothesis (A2). (A1) and (A2) are not mutually exclusive, as sensory regions may represent both stimulus features and environmentally relevant feature conjunctions.

By “perceived emotions” we refer only to participants’ perceptions of the stimuli and their judgments of what emotions the stimuli expressed. We do not refer to emotional states evoked in the participants by the stimuli, or to any other kind of emotion. Importantly, though we discuss the relevance of the findings to cross-cultural generalization, we did not test any hypotheses across cultures.

Testing both sets of hypotheses required comparing representations between brain areas. To accomplish this, we used model-based representational similarity analysis (RSA) (Kriegeskorte, Goebel, & Bandettini, 2006; Kriegeskorte, Mur, & Bandettini, 2008), comparing representations evoked by separately presented auditory and visual stimuli to test (H1) and (H2). For detailed discussion of the limits and merits of this approach, see Roskies (2021). The model included predictors corresponding to both simple stimulus features and to participants’ judgments of emotion content, supporting tests of (A1) and (A2). We performed an additional supporting test of (H1) using a model-free approach that directly compared representational geometries across sensory areas without making any assumptions about representational content.

### Previous research on neural representation of emotion

Emotion-related neural processes are distributed across a wide range of brain areas, with each area implicated in the production and/or perception of many emotions (Lindquist, Wager, Kober, Bliss-Moreau, & Barrett, 2012; Wager et al., 2015). However, certain aspects of emotion processing are localized. Lesion and neuroimaging studies have demonstrated that some brain areas play an outsized role in the processing of specific emotions; for example, the amygdala for the conscious recognition of fearful stimuli (Adolphs, Tranel, Damasio, & Damasio, 1994; Tsuchiya, Moradi, Felsen, Yamazaki, & Adolphs, 2009), and the insula for recognizing disgust (Calder, Lawrence, & Young, 2001; Phillips et al., 1997). Because our hypotheses concern representations capable of distinguishing many different emotion expressions, we focus here on distributed representations of emotion, and not on areas implicated in processing individual emotions.

Our hypotheses ask not only *where* in the brain emotions are represented, but *how* those representations are structured. For example, a single brain area may distinguish stimulus classes using different spatial patterns of activity that all have the same mean. To characterize the representational properties of these areas, it is necessary to use techniques that are sensitive to such spatially distributed patterns; e.g., multivariate pattern classification (Norman, Polyn, Detre, & Haxby, 2006) or representational similarity analysis (RSA; Kriegeskorte & Kievit, 2013). For example, Peelen et al. (2010) showed that medial prefrontal cortex (mPFC) and posterior superior temporal sulcus (pSTS) supramodally represent emotion identity by demonstrating that patterns of activity in these areas had greater within-emotion similarity than between-emotion similarity. Chikazoe et al. (2014) used pattern analysis to locate supramodal valence (positive vs. neutral vs. negative) representations in medial and lateral orbitofrontal cortex and modality-specific valence representations in sensory cortices. Also investigating valence, Kim et al. (2017) presented emotional movie clips and orchestral music, and found a range of supramodal representations: valence direction in the precuneus, valence magnitude in mPFC, STS, and middle frontal gyrus (MFG), and both valence direction and magnitude in the STS, MFG, and thalamus. Skerry & Saxe (2015) found that a model describing participants’ appraisals of emotional narratives (e.g., “Did someone cause this situation intentionally, or did it occur by accident?”) fit activity in dorsal and middle medial prefrontal cortex, the temporoparietal junction, and a network of regions identified by a theory of mind localization task.

Importantly, where previous studies have focused on emotions evoked by narrative content while controlling for stimulus features (e.g., Chikazoe et al., 2014; Kim et al., 2017; Skerry & Saxe, 2015), the present study takes a different approach, focusing on emotion perceived solely from stimulus features without any contextualizing narrative. Emotions perceived from stimulus features make up a large and understudied part of human experience. For example, people often communicate emotion using only body language and tone of voice, and actively seek out instrumental music and abstract visual art that communicates emotion only through variation in pitch, volume, shape, brightness, and so on. And although there are many cross-cultural differences in emotion experience, expression, and perception (Gendron et al., 2014; Jack et al., 2012, 2016; Jackson et al., 2019; Margulis et al., 2019; Yuki et al., 2007), preliminary evidence suggests the use of shared stimulus features to express emotion can generalize across dissimilar cultures (Sievers et al., 2013; Trehub et al., 2015). Despite its ubiquity and importance, the neural mechanisms supporting emotion perception from stimulus features remain poorly understood.

The present approach allows us to test the *shared features* (A1) and *environmental conjunctions* (A2) hypotheses, assessing whether sensory brain areas represent conjunctions of features associated with environmentally relevant stimuli such as emotion expressions, or whether these areas represent simple features that may be used by separate, downstream areas to infer emotion content.

### Stimuli and experimental paradigm

The stimuli consisted of short piano melodies and animations of a bouncing ball generated by participants in a previous study. This study showed that emotions were expressed the same way in music and in movement in both the US and a small-scale society in rural Cambodia (Sievers et al., 2013). The participants used a computer program to create examples of five emotions (Angry, Happy, Peaceful, Sad, Scared) by manipulating five stimulus features (speed, irregularity, consonance/spikiness, ratio of big-to-small movements, ratio of upward-to-downward movements). Participants were split into separate music and movement groups, each of which had no knowledge of the other. This approach did not presuppose what combinations of features would be used for each emotion, and participants were not instructed to use any specific features or feature combinations. Instead, they were encouraged to explore the entire possibility space. Critically, this method allowed us to vary what emotions were communicated while holding the depicted objects constant (i.e., each emotion was communicated using only the piano or the bouncing ball). This guaranteed that emotion content could only be communicated by variation in stimulus features, and that processing requirements were consistent across the stimulus set.

Note that this approach differs from previous research where emotion was communicated using narrative stories or emotionally charged images; e.g., the International Affective Picture System (Lang, Bradley, & Cuthbert, 2008). Such studies often control for stimulus features, guaranteeing that emotion judgments are based solely on the content depicted in the stimuli. For example, a study of perceived emotion in spoken narrative might control for the speaker’s tone of voice, focusing on what the speaker said, rather than how they said it. The present study takes the opposite approach, controlling for the content depicted in the stimuli, guaranteeing that participants’ emotion judgments are based solely on variation in stimulus features. This is analogous to holding a speaker’s words constant, so that emotion can only be communicated by tone of voice.

Because many emotions are perceived as mixes of other emotions (Cowen & Keltner, 2017), the stimulus set was augmented by linearly mixing the features of each emotion pair, creating mixed emotions (e.g. Happy–Sad). Emotions were mixed at 25%, 50%, and 75%. Three additional, “neutral” emotions were identified by searching for points in the stimulus feature possibility space that were distant from all other emotional feature combinations. For each set of stimulus features, or stimulus class, many individual stimuli were probabilistically generated (see *Detailed methods*). This ensured the results were not dependent on the idiosyncracies of single stimuli, but were instead generalizable to all stimuli that shared the same features. Further, this prevented participants from memorizing arbitrary associations between individual stimuli and emotion labels. Music and animation were matched, such that for each musical stimulus class there was an animation stimulus class with the same features. This process yielded 76 total emotional stimulus classes, including both music and animation. All stimuli are available at https://osf.io/kvbqm/.

A separate set of participants judged how well each stimulus fit all five emotion labels, and a subset of these participants viewed many music and animation stimuli while undergoing functional magnetic resonance imaging (fMRI) (Figure 1).

**Figure 1:**
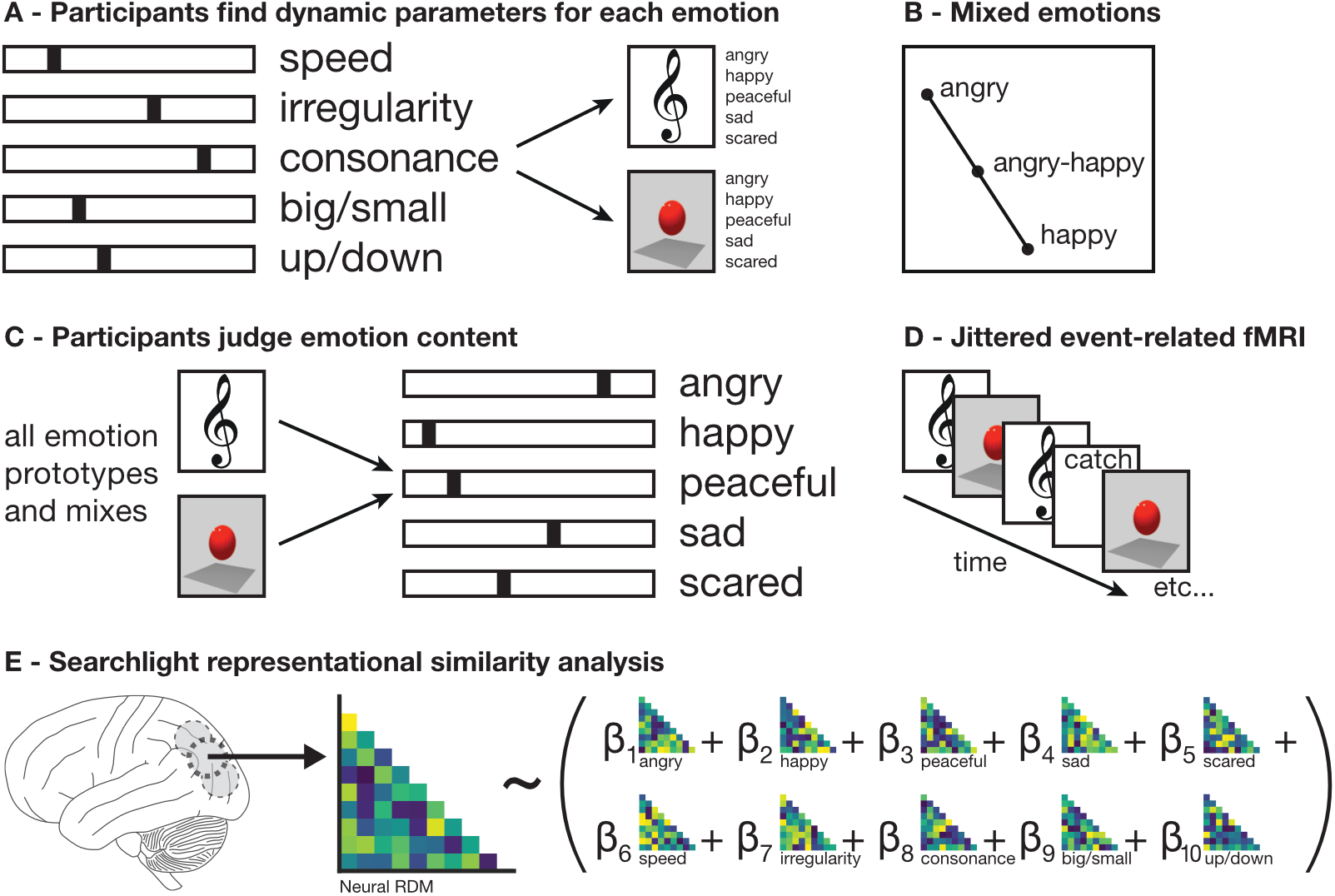
Experimental paradigm. *A*. Participants in Sievers et al. (2013) manipulated stimulus features to generate music and animation communicating five prototypical emotions: Angry, Happy, Peaceful, Sad, and Scared. *B*. Mixed emotions were generated by linear interpolation between the stimulus features of prototypical emotions. *C*. Participants judged the emotion content of many prototypical and mixed emotions in music and animation. *D*. A subset of participants viewed many prototypical and mixed emotions in music and animation while undergoing jittered event-related fMRI scanning. *E*. Results were analyzed using searchlight representational similarity analysis (Kriegeskorte et al., 2006, 2008; Kriegeskorte & Kievit, 2013). For each searchlight sphere, the structure of the neural representational dissimilarity matrix (RDM) was predicted using a linear combination of stimulus feature and emotion judgment RDMs.

## Results

### Emotion judgments

Participants broadly agreed about the emotion content of each stimulus class (Figure 2). Agreement was assessed by measuring the distance of participants’ individual emotion judgments from the class mean, scaled by the maximum possible distance, and significance was assessed using permutation testing (see *Detailed methods*). For all 76 stimulus classes except one “neutral” emotion, participants’ judgments were closer to the class mean than would be expected by chance (mean *t*=−4.37; mean difference from null=.07; mean p<.001). Importantly, this agreement rules out the possibility that participants invented and then memorized arbitrary associations between combinations of stimulus features and combinations of emotion labels.

**Figure 2:**
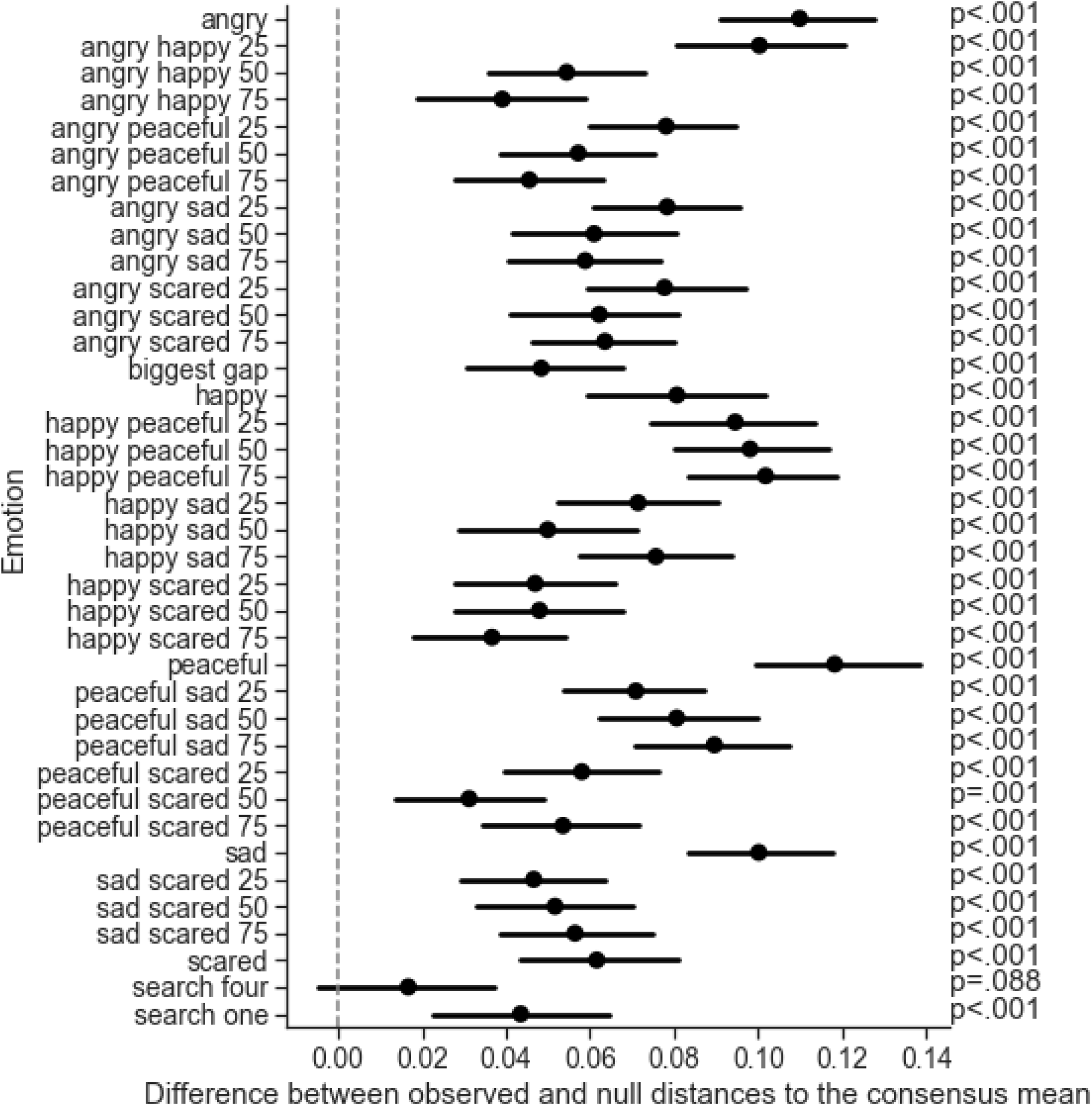
Emotion judgment agreement. Agreement between participants was assessed by measuring the distance of participants’ individual emotion judgments from the class mean. Significance was assessed using permutation testing (see *Detailed methods*). Values above 0 indicate more agreement than expected by chance.

### Shared representational geometry

Auditory and visual brain regions shared a representational geometry. A single model of representational similarity (Kriegeskorte et al., 2006, 2008; Kriegeskorte & Kievit, 2013) explained patterns of activity in visual brain regions during animation trials and auditory brain regions during music trials, providing strong support for the *separate regions, shared representations* hypothesis (H1) (Figure 3; Table 1). The model used 10 representational dissimilarity matrices (RDMs) as predictors: five based on the mean parameter settings used to create the stimuli (speed, irregularity/jitter, consonance/spikiness, ratio of big-to-small movements, ratio of upward-to-downward movements), and five based on the mean emotion judgments of the behavioral participants (Angry, Happy, Peaceful, Sad, and Scared) (Figure 4). The model included no information specific to either vision or audition.

**Figure 3:**
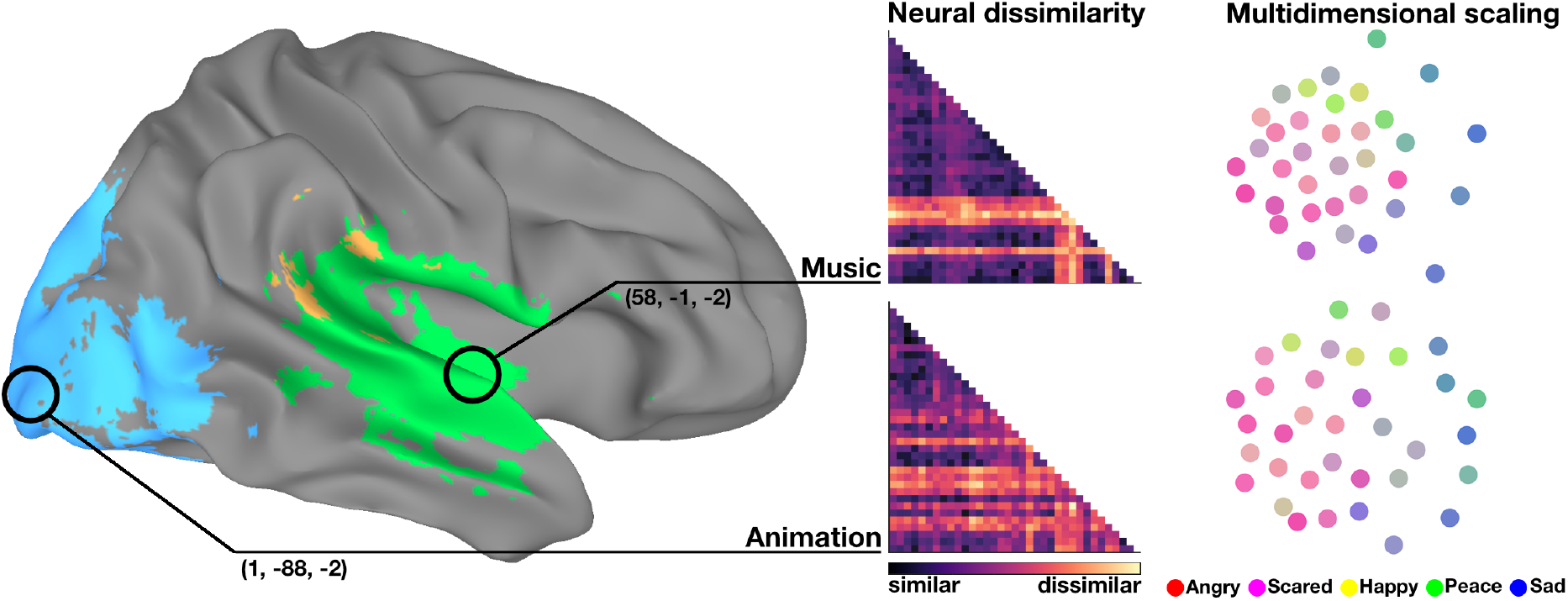
Main result. Highlighted brain areas were identified using a model including both stimulus features and emotion judgments as predictors, which was separately fit to animation trials (blue) and music trials (green). A significant proportion of participants’ model fits overlapped for both trial types (yellow). Neural dissimilarity matrices show pairwise distances between activity patterns evoked by each stimulus at the locations of best model fit (circled)—medial lingual gyrus (animation) and lateral superior temporal gyrus (music). Labels as in Figure 4. Multidimensional scaling flattens these matrices to two dimensions, so the distance between dots reflects the similarity of patterns of neural activity. Dots are colored by mixing the legend colors based on participants’ judgments of the emotion content of each stimulus.

**Table 1:** Peak model fits. Anim.: Model fit to animation trials. Music: Model fit to music trials. Overlap: Percentage of participants with overlapping music and animation model fits. Inter.: Intermodal regions which fit the model even when the stimulus was presented in the non-preferred modality. For results per model predictor, see Figures S4 and S5. Labels determined programmatically using the atlas of Destrieux, Fischl, Dale, and Halgren (2010).

| Analysis | x, y, z | Nearest atlas label (2010) | $R^2_{adj}$ | % | 95% CI | p |
| --- | --- | --- | --- | --- | --- | --- |
| Music | 58, -2, -2 | R Lateral aspect of the superior temporal gyrus | .15 |  | .10–.20 | .011 |
| Music | -62, -16, 7 | L Lateral aspect of the superior temporal gyrus | .09 |  | .05–.12 | .011 |
| Anim. | 2, -88, -2 | L Lingual gyrus, lingual part of the medial occipito-temporal gyrus, (O5) | .15 |  | .08–.21 | .005 |
| Anim. | 46, -68, 1 | R Inferior occipital gyrus (O3) and sulcus | .04 |  | .01–.07 | .005 |
| Anim. | 22, -82, 31 | R Superior occipital gyrus (O1) | .03 |  | .01–.06 | .005 |
| Overlap | 64, -28, 22 | R Supramarginal gyrus |  | 60% | .36–.84 | <.001 |

| Analysis | x, y, z | Nearest atlas label (2010) | $R_{adj}^2$ | % | 95% CI | p |
| --- | --- | --- | --- | --- | --- | --- |
| Overlap | -58, -34, 19 | L Supramarginal gyrus |  | 40% | .16–.64 | .008 |
| Inter. | 2, -88, -2 | L Lingual gyrus, lingual part of the medial occipito-temporal gyrus, (O5) | .28 |  | .20–.37 | <.001 |
| Inter. | 64, -28, 22 | R Supramarginal gyrus | .09 |  | .06–.12 | <.001 |
| Inter. | -56, -40, 22 | L Planum temporale or temporal plane of the superior temporal gyrus | .08 |  | .05–.10 | <.001 |
| Inter. | 32, -56, 61 | R Superior parietal lobule (lateral part of P1) | .07 |  | .05–.09 | <.001 |
| Inter. | -32, -56, 64 | L Superior parietal lobule (lateral part of P1) | .06 |  | .04–.08 | <.001 |
| Inter. | -16, -22, 40 | L Marginal branch (or part) of the cingulate sulcus | .05 |  | .03–.07 | <.001 |
| Inter. | -28, -58, -53 | L Lateral occipito-temporal gyrus (fusiform gyrus, O4-T4) | .04 |  | .03–.06 | <.001 |
| Inter. | -46, 44, 22 | L Middle frontal gyrus (F2) | .03 |  | .02–.04 | <.001 |
| Inter. | -4, 64, 22 | L Superior frontal gyrus (F1) | .03 |  | .02–.04 | <.001 |

**Figure 4:**
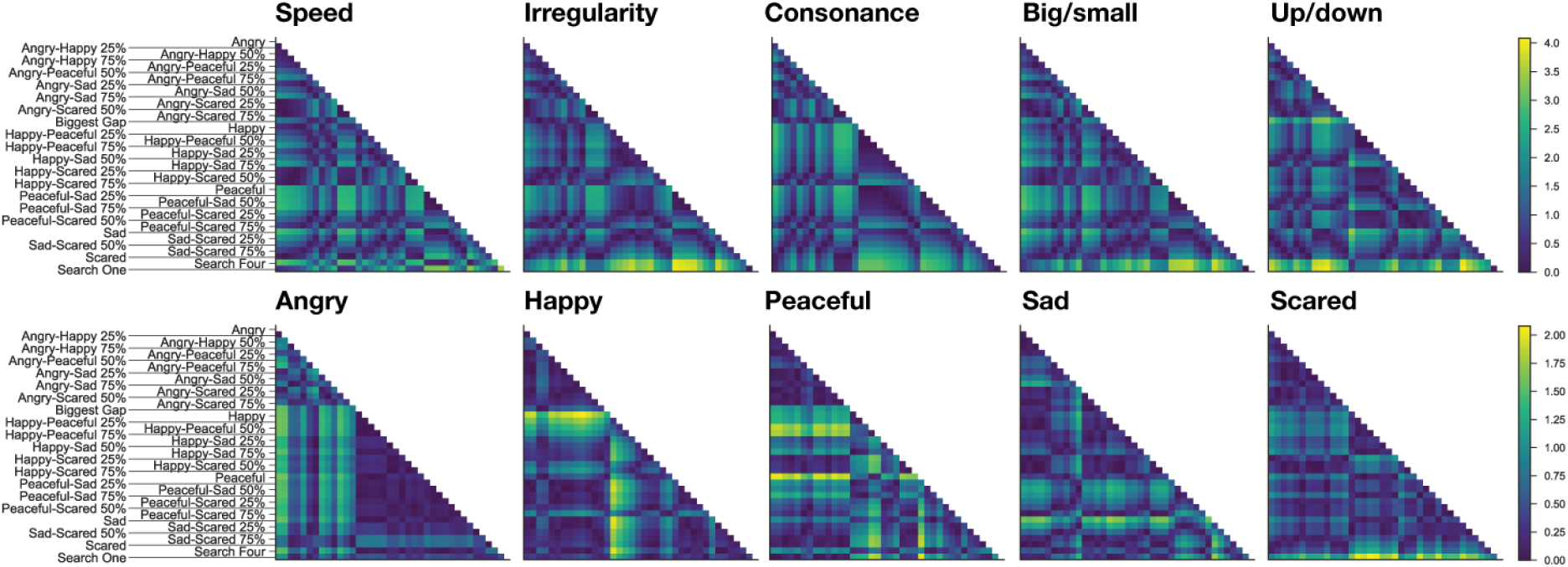
Representational dissimilarity matrices. Columns and rows share labels. “Biggest Gap,” “Search One”, and “Search Four” are “neutral” emotions.

The peak of the average model fit across participants was in the left medial lingual gyrus for animation trials (mean 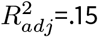; 95% CI: .08–.21; t(19)=4.68; p=.005 corrected) and in right anterior superior temporal gyrus for music trials (mean 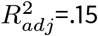; 95% CI: .1–.2; t(19)=6.08; p=.01 corrected), (Figure 3). Critically, a direct, model-free test of similarity between these areas showed that they were more similar to each other than would be expected by chance (ρ=.68, p<.001), further supporting the *separate regions, shared representations* hypothesis (H1), and making it unlikely that the results reported above are an artifact of model misspecification (see *Detailed methods*).

Model fit was driven by both stimulus feature *and* emotion judgment predictors, and was not dominated by a small number of predictors, providing support for both the *simple features* and *environmental conjunctions* hypotheses (A1 and A2). Individual predictors were assessed by mapping Spearman’s ρ across the brain. Spearman’s ρ was significant for all 10 predictors at the location of peak model fit (Figure 5), and was distributed similarly across the brain (Figure S4). β weight maps for each predictor were also calculated (Figure S5), reflecting only the unique contribution of each predictor, whereas Spearman’s ρ reflects both unique and shared contributions. See Figure S6 for an assessment of model multicollinearity, including variance inflation factors for each predictor.

**Figure 5:**
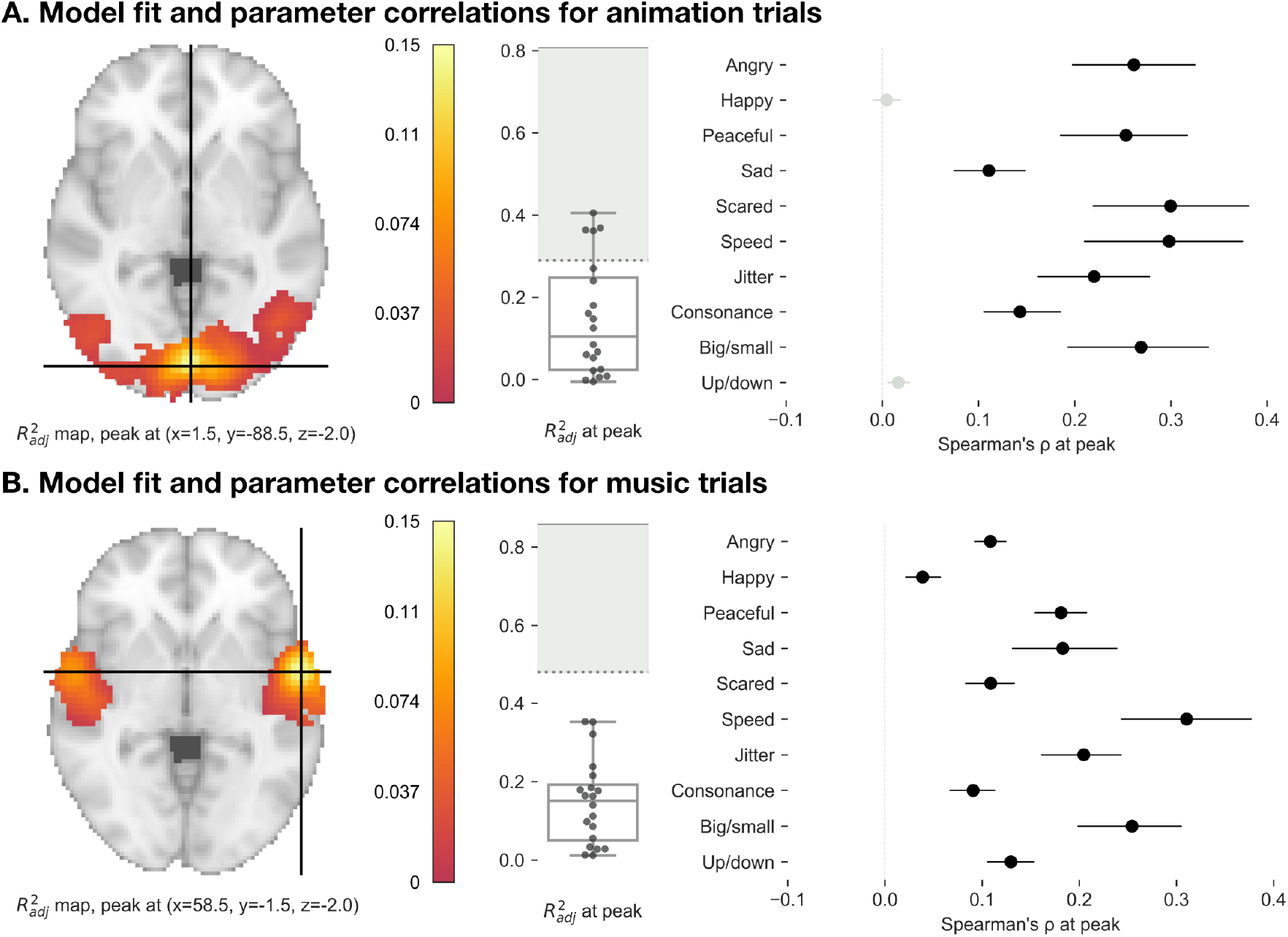
Model fits. Maps of the mean coefficient of determination 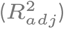 across participants. Maps thresholded at FWER=.05. Box plots show the median, quartiles, and range of the per-participant 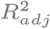 values at the location of best model fit at the group level. The dotted line indicates the lower bound of the noise ceiling, and the solid line the upper bound. For per-parameter Spearman’s ρ and β weight maps, see Figures S4 and S5.

The model accounted for 51% of the variance for animation trials, and 31% of the variance for music trials, relative to the lower bound of the noise ceiling (see *Detailed methods*). Note that because of small differences in functional anatomy across participants, the peak of the average model fit underestimates individual model fit. The mean of the individual peak model fits was in bilateral anterior superior temporal gyrus for music trials (mean individual 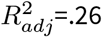; 95% CI: .21–.31; t(19)=10.95; p<.001 uncorrected) and in the lingual gyrus for animation trials (mean individual 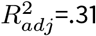; 95% CI: .24–.38; t(19)=9.2; p<.001 uncorrected) (Figures S1 and S2).

### Overlapping auditory and visual model fit

Brain regions where music and animation were both represented were found in bilateral posterior superior temporal gyrus (pSTG) in 60% of participants (95% CI 36%–84%, p<.001 corrected), supporting the *supramodal region* hypothesis (H2) (Figure 6A; see Figure S3 for per-participant maps). To locate such supramodal representations we created binary overlap masks, selecting voxels where both music and animation model fits were significant at the individual level (permutation p<.05 uncorrected). Multiple comparisons correction of these overlap maps was performed at the group level, testing the proportion of individuals with overlap in a region against the null hypothesis that no participants had overlap in that region. Critically, this analysis is insensitive to the magnitude of 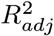 at the individual level, allowing detection of overlapping signals that have low magnitude but reappear across a significant proportion of the participants. The model fit for music trials was also significant at this location, though the model fit for animation trials was not (music mean 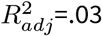, 95% CI: .02–.05, t(19)=5.78, p=.01 corrected; animation mean 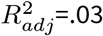, 95% CI: .01–.05, t(19)=2.94, p=.13 corrected). The model accounted for 26% of the variance for animation trials, and 31% of the variance for music trials, relative to the lower bound of the noise ceiling. Due to individual differences in functional anatomy, this procedure underestimates the proportion of participants with supramodal representations.

**Figure 6:**
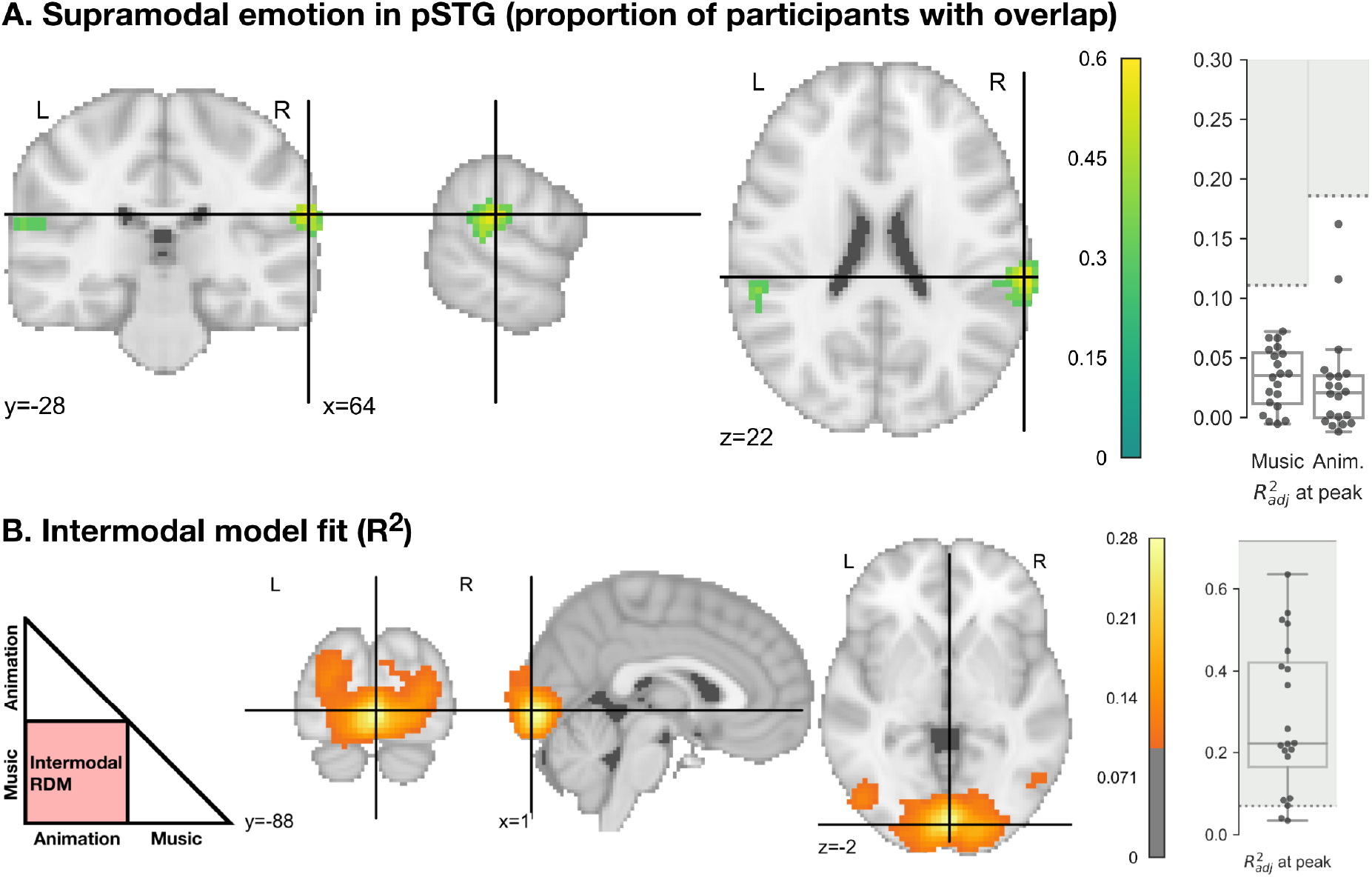
Results across modalities. A. Supramodal emotion in pSTG. Maps show the proportion of participants representing emotion in music and animation in the same brain areas, thresholded at voxelwise FWER=.05. Box plots show the median, quartiles, and range of 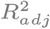 for music and animation trials at the marked peak. B. Intermodal RSA model fit. Maps show areas that represented emotional stimuli even when presented in the area’s non-preferred modality (see *Detailed methods*), thresholded at voxelwise FWER=.05. 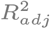 values below .1 hidden for visual clarity. Box plot shows the median, quartiles, and range of per-participant 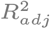 values at the marked peak. Dotted lines indicate the lower bound of the noise ceiling, while solid lines indicate the upper bound.

### Exploratory intermodal RSA

To find brain areas that represented stimuli presented in that area’s non-preferred modality, we performed an exploratory intermodal RSA (see *Detailed methods*). Intermodal RSA revealed a bilateral set of areas across occipital, superior parietal, temporal, cingulate, and frontal cortex that represented stimuli presented in their non-preferred modality (Figure 6B; Table 1). Note that some of these areas did not show significant unimodal model fit. Peak intermodal model fit was in left lingual gyrus (mean 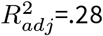; 95% CI: .20–.37; t(19)=6.9; p<.001 corrected). Notably, the peak intermodal model fit exceeded the peak within-modality model fit for both music and animation, and also exceeded the lower bound of the noise ceiling, explaining 40% of the variance relative to the upper bound. This suggests intermodal activity in left lingual gyrus was dominated by representations of model features. However, the lower bound of the intermodal noise ceiling was relatively low (.07), suggesting that most reliable neural activity in this region was modality-specific.

## Discussion

Music and movement are subjectively linked, and both use similar features to communicate emotion content. We examined a possible explanation for this link: that the brain represents music and movement using a shared representational geometry. To investigate this, we tested two primary hypotheses. (H1) The *separate regions, shared representations* hypothesis, where separate auditory and visual regions use the same representational geometry; and (H2) the *supramodal region* hypothesis, where some region(s) represent both auditory and visual stimuli. We also tested two auxiliary hypotheses. (A1) The *simple features* hypothesis, where sensory areas represent individual stimulus features that are not directly associated with emotion content; and (A2) the *environmental conjunctions* hypothesis, where sensory areas represent conjunctions of features that directly track differences in emotion judgment.

We found that brain activity in separate auditory and visual areas shared a representational geometry, supporting the *separate regions, shared representations* hypothesis (H1). Providing additional support for (H1), representations in auditory and visual brain areas were more similar to each other than to any randomly chosen pair of brain areas. Further, music and animation were represented in pSTG, suggesting the pSTG uses a supramodal representation, supporting the *supramodal region* hypothesis (H2).

Stimulus feature predictors (speed, jitter, consonance/spikiness, ratio of upward-to-downward movements, and ratio of big-to-small movements) were significant in both auditory and visual regions, supporting the *simple features* hypothesis (A1). A shared, crossmodal representation of simple stimulus features would support downstream comparison of auditory and visual stimuli, including inferential assessment of emotion content by, e.g., simulation theory (Gordon, 1986) or theory theory (Gopnik & Wellman, 1994) systems. On such an account there may be nothing emotional *per se* about representations in sensory brain regions. However, predictors based on participants’ emotion judgments were also significant in both auditory and visual regions, even when controlling for the stimulus feature predictors, supporting the *environmental conjunctions* hypothesis (A2). On this account, sensory regions represent conjunctions of task-relevant environmental features, such as those associated with emotion expressions, supporting direct perception of social information (Chemero, 2006; Gallagher, 2008).

Other possible parameters such as valence and arousal (Russell, 1980), value (Levy & Glimcher, 2012; Shuster & Levy, 2018), Fourier features (Sievers et al., 2019), HMAX (Riesenhuber & Poggio, 1999), and motion energy (Nishimoto et al., 2011) certainly covary with the stimulus feature and emotion judgment predictors. Because these and similar measures would be fully dependent on the model parameters, including them as controls would introduce collinearity and create post-treatment bias. Identifying exactly how the features used in the reported model map to the true dimensions on which emotion, audition, and vision are organized will require future research.

An exploratory intermodal representational similarity analysis found that visual areas represented both stimulus feature and emotion judgment predictors when musical stimuli were presented. However, most reliable neural activity in these areas was modality-specific, as indicated by a low intermodal noise ceiling. Previous studies have shown multimodal processing in unimodal areas (for reviews, see Bulkin & Groh, 2006; Ghazanfar & Schroeder, 2006; Kayser & Logothetis, 2007), which may depend on projections between unimodal areas (Cappe & Barone, 2005; Falchier, Clavagnier, Barone, & Kennedy, 2002; Rockland & Ojima, 2003). The reported results extend this account by showing that crossmodal perception is the product not only of operations in association cortices or activity dependent on inter-areal projections, but of the use of a representational geometry that is shared across modalities.

The reported findings in pSTG are near previously reported pSTS activation during action understanding (Beauchamp, Lee, Argall, & Martin, 2004; Wyk, Hudac, Carter, Sobel, & Pelphrey, 2009), emotion perception (Kreifelts, Ethofer, Grodd, Erb, & Wildgruber, 2007; Robins, Hunyadi, & Schultz, 2009; Watson et al., 2014), affective and linguistic prosody recognition (Belyk & Brown, 2014), and crossmodal perception and recognition tasks (Werner & Noppeney, 2010; Wright, Pelphrey, Allison, McKeown, & McCarthy, 2003). Interestingly, the reported results were right lateralized, similar to previous findings on prosody recognition (Belyk & Brown, 2014). Damage to the pSTS does not impair voice recognition (Jiahui et al., 2017), suggesting its representations are downstream from feature detectors. Alongside these results, the reported findings are consistent with the hypothesis that the pSTG/pSTS acts as a hub for transforming unimodal inputs into a common supramodal representation (Schirmer & Adolphs, 2017).

### Evoked emotion

Although our participants perceived emotions in our stimuli, it is unlikely that our stimuli evoked emotions in our participants. This disjunction highlights the complex and sometimes paradoxical relationship between perceived and evoked emotion. For example, perceiving sadness in music can evoke feelings of romance or pleasure (Kawakami, Furukawa, Katahira, & Okanoya, 2013). The gap between perception and feeling has been theorized in terms of direct versus vicarious emotions (Kawakami et al., 2013), and in terms of emotion modules serving complementary functions (Gelstein et al., 2011; Mehr, Krasnow, Bryant, & Hagen, 2020). Another possibility is that perceptual representations of stimulus features and emotion content interact with regions that produce context-sensitive appraisals and emotion experiences, such as those identified by Skerry & Saxe (2015), and subcortical regions sensitive to emotion content, including the amygdala (Wang et al., 2014). Activation of these appraisal and experience-related regions may not be necessary for making simple judgments of emotion content from stimulus features, possibly accounting for their absence in our results.

However, perceptual representations of emotion may also be linked to evoked emotions. Saarimäki et al. (2018) showed that emotions evoked by listening to short stories produced activity in visual cortex, suggesting that evoked emotions can activate associated sensory representations. This may be a special case of the more general principle that mental imagery and episodic memory depend in part on activity in sensory regions associated with similar experiences (Wheeler, Petersen, & Buckner, 2000). Accordingly, perceptual representations of emotion content may form over development in a process similar to memory consolidation. This developmental process may be guided by language, supporting culture-specific particularity (Barrett, Lindquist, & Gendron, 2007; Hoemann, Xu, & Barrett, 2019). Activation of perceptual representations of emotion by imagined emotion experience could play an important role in art and music by allowing artists and composers to iteratively check whether their artistic products correspond with their perceptual representations.

### Systematicity, iconicity, and conceptual scope

The neural representational system identified here is likely involved in phenomena beyond emotion perception, raising an interesting question: What concepts can and cannot be communicated via combinations of crossmodal stimulus features? If feature combinations in music and movement are symbolic, like words in natural language, then we would expect stimulus feature combinations that refer to abstract, non-emotional concepts. Just as arbitrary sequences of phonemes can point to “the housing market” or “editorial policy,” arbitrary combinations of stimulus features should be able to do the same.

But, strikingly, music and movement do not operate wholly on an arbitrary, symbolic basis. Music and movement systematically use variation in the magnitudes of stimulus features to communicate variation in the magnitudes of concepts which the stimulus iconically resembles (Sievers et al., 2019, 2013; Spector & Maurer, 2009). For example, participants in the present study perceived mixes of the features for “happy” and the features for “sad” as expressing emotions on a continuum between happiness and sadness, with this pattern generalizing across emotion pairs. The present results suggest this systematic mixing is made possible in part by a crossmodally shared neural representational geometry.

The systematic and iconic properties of musical communication may partially account for its use in expressing emotion, even across different cultural contexts. For example, the stimulus generator for the present study was previously used to show that the same combinations of music and movement features express the same emotions in both the United States and a small-scale society in rural Cambodia (Sievers et al., 2013). It may be that the forms of emotional music and movement are fixed by iconic, functional relationships that are shared across cultures (Mehr et al., 2020, p. @Sievers2021). This may be why lullabies are slow and consonant across a global sample of ethnographic reports and recordings (Mehr et al., 2019), and why high emotional arousal is expressed using harsh sounding, high spectral centroid sounds even across species of terrestrial vertebrates (Filippi et al., 2017).

Music, movement, and crossmodal neural representations may be well-suited to communicate iconic concepts that vary in magnitude, whereas language may be well-suited to communicate symbolic concepts that vary in kind. Similarly, it may be that iconic communication tends to generalize across cultures while symbolic communication tends to be more culture-specific. For example, previous research has shown cross-cultural generalization of valence perception but not categorical emotion (Gendron et al., 2014) (though categorical emotions can also be shared across cultures (Parkinson, Walker, Memmi, & Wheatley, 2017)). Further, musical narrative built from contrasting sets of stimulus features (Margulis, 2017) shows large cross-cultural variability in interpretation (Margulis et al., 2019), unlike the present stimuli (Sievers et al., 2013) which contained no such contrasts.

Importantly, the present study did not test any hypotheses across cultures. Future research will need to explore broad areas of concept space across many cultures, collect free responses from participants, characterize culture-specific emotion concepts, and contrast concepts that vary in magnitude with concepts that vary in kind.

### Direct perception

The results support the *environmental conjunctions* hypothesis (A2), that sensory brain regions represent task-relevant combinations of stimulus features, reducing the need for downstream inferential processing and acting as a shortcut for making important judgments. These representations may provide a neural basis for the direct perception of social information (Chemero, 2006; Gallagher, 2008)— exemplified here by emotion judgment, and potentially covering a range of other phenomena. Importantly, the *simple features* hypothesis (A1) was also supported, suggesting that direct perception and inferential processing systems coexist and may interact.

Without contextualizing narrative, judgments of emotion content in music and movement depend on configurations of stimulus features (Sievers et al., 2019, 2013) in much the same way that the solution to a puzzle depends on the configuration of the pieces. In other words, stimulus features and emotion judgments are naturally confounded. The crux of the *environmental conjunctions* hypothesis (A2) is that any combination of features that is sufficiently confounded with a target is useful for identifying that target. We argue that the brain uses such natural confounds as a shortcut to make task-relevant judgments: if sensory regions represent feature combinations that are perfectly confounded with a target’s identity, downstream inferential processing may not be necessary to identify the target.

In demonstrating support for the *environmental conjunctions* hypothesis (A2), we do not mean to suggest that sensory brain areas alone support purely conceptual, symbolic, or cognitive labeling of emotions. Previous studies using context-dependent and narrative stimuli have demonstrated the importance of inferential processing for emotion perception (Barrett, Mesquita, & Gendron, 2011; Skerry & Saxe, 2015). Further, inferential processing may play a role in the gradual tuning of perceptual systems for direct perception across development. Simple adaptations for perceiving cross-sensory magnitude or position information (Deneux et al., 2019; Murphy et al., 2020) or for adaptive signalling (Hebets et al., 2016; Huron, 2012; Johnstone, 1996, 1997) may work in concert with learning (Clark, 2013; Kok, Brouwer, Gerven, & Lange, 2013; Lange, Heilbron, & Kok, 2018; Saffran, Aslin, & Newport, 1996), language (Hoemann et al., 2019), and cultural evolution (Laland, Odling-Smee, & Feldman, 2000) processes to support the development of task-relevant representations. This arrangement could flexibly accommodate culture-specific emotion concepts and display rules (Jack et al., 2012, 2016; Yuki et al., 2007).

Such tuning of sensory representations to the features used to communicate and categorize emotions shows that the need to identify such signals has exerted a profound shaping force on perceptual processes. We do not see or hear the actions of others as raw sense impressions first, later decode their conceptual content, and finally make an abstract emotion judgment. Rather, we begin accumulating evidence for emotion judgments from the lowest levels of sensory processing.

## Acknowledgements

We thank Sam Nasatase, Matteo Visconti di Oleggio Castello, J. Swaroop Guntupalli, and Joshua Greene for helpful comments during the writing process, and Paulina Calcaterra, Rebecca Drapkin, Caitlyn Lee, Elizabeth Reynolds, Tshibambe Nathanael Tshimbombu, and Kelsey Wheeler for assistance collecting fMRI data. This research was supported in part by the Nelson A. Rockefeller Center for Public Policy at Dartmouth, the John Templeton Foundation, the Neukom Institute for Computational Science, the Vision Science to Applications (VISTA) program funded by the Canada First Research Excellence Fund (CFREF, 2016–2023) and by the Natural Sciences and Engineering Research Council of Canada.

## Author contributions

B. Sievers: Conceptualization, data curation, formal analysis, funding acquisition, investigation, methodology, project administration, software, vizualization, writing - original draft, review & editing. C. Parkinson: Investigation, methodology, writing - review & editing. P.J. Kohler: Methodology, writing - review & editing. J.M. Hughes: Methodology, writing - review & editing. S.V. Fogelson: Methodology, writing - review & editing. T. Wheatley: Conceptualization, funding acquisition, resources, supervision, writing - review & editing. Roles defined by the Contributor Roles Taxonomy, available at https://casrai.org/credit/

## Detailed methods

### Lead contact

Further information and requests for resources should be directed to and will be fulfilled by the lead contact, Thalia Wheatley.

### Materials availability

Stimuli have been deposited to osf.io (Sievers, 2021).

### Data and code availability

De-identified fMRI data have been deposited to OpenNeuro (Sievers et al., 2021). All original code has been deposited to osf.io (Sievers, 2021). Any additional information required to reanalyze the data reported in this paper is available from the lead contact upon request.

### Experimental model and subject details

79 participants (47 female) were recruited from the Dartmouth College student community to participate in the emotion evaluation task (experiment 1). 20 of these participants (11 female) also participated in the fMRI of emotion viewing task (experiment 2). All fMRI participants were right-handed and had normal or corrected-to-normal vision. All participants provided written informed consent, and the study was approved by the Dartmouth College Committee for the Protection of Human Subjects.

### Method details

#### Stimuli

Emotion stimuli were generated using a model developed for a prior study (Sievers et al., 2013) that used movement across a number line to create both music (simple piano melodies) and animated movement (a bouncing ball). The model had five stimulus feature parameters: speed, irregularity/jitter, consonance/spikiness, ratio of big-to-small movements, and ratio of upward-to-downward movements. Each time the model was run, it probabilistically generated a new stimulus based on its current parameter settings. Participants in Sievers et al. (2013) (music *N*=25; movement *N*=25; total *N*=50) used this model to communicate five prototype emotions: Angry, Happy, Peaceful, Sad, and Scared. Critically, participants were split into separate music and movement groups, each of which had no knowledge of the other. Participants chose similar music and movement parameter settings for each emotion across modalities, showing that music and movement share an underlying structure. The median parameter settings across music and movement from the United States participants in Sievers et al. (2013) were used to generate the stimuli used in the present studies. All stimuli are available at https://osf.io/kvbqm/.

In addition to the prototype emotions, mixed emotion stimuli were created by interpolating linearly between the parameter settings for each prototype emotion pair; 25%, 50%, and 75% mixes were used. We also added three putatively “neutral” or “non-emotional” parameter settings that were selected to be distant from all other stimuli. “Search One” and “Search Four” were selected by a Monte Carlo search algorithm, and consisted of extreme values for all five parameters. “Biggest Gap” was created by selecting the midpoint of the largest gap between the five prototype emotions and the stimulus feature parameter endpoints.

For each prototype, mixed, and “non-emotional” parameter setting in each modality, we probabilistically generated 20 exemplars, for a total of 1,520 stimuli (38 emotions x 2 modalities x 20 exemplars). To eliminate the possibility of generating unusual outlier stimuli, each candidate exemplar was compared to a larger, separate sample of 5000 same-emotion exemplars, and was re-generated if found to be further than one standard deviation from the emotion mean along any parameter.

#### Experiment 1 (emotion evaluation)

Participants (*N*=79, 47 female) evaluated the emotion content of the stimuli. Stimuli were presented using a computer program developed using Max/MSP version 5 (Zicarelli, 1998) that displayed five slider bars, one for each emotion prototype (Angry, Happy, Peaceful, Sad, and Scared). The on-screen order of slider bars and emotion stimuli were randomized across participants. Participants viewed or listened to each stimulus at least three times, and were asked “to evaluate the amount or intensity of emotion expressed by the music or animation by positioning the slider bars.”

#### Experiment 2 (fMRI of emotion viewing)

During each fMRI run, participants (*N*=20, 11 female) viewed 18 randomly selected exemplars from each of the 76 stimulus classes described above. Each stimulus class was shown once per run, and participants completed 18 runs across 3 separate scanning sessions (~3 hours of scan time, 1,368 stimulus impressions). Each scan session was scheduled for approximately the same time of day, and no more than one week elapsed between scan sessions.

Stimuli were truncated to 3s in duration and followed by fixation periods of randomly varying duration (range: 0.5s–20s). The ratio of stimulus presentation to fixation was 1:1. A Monte Carlo procedure was used to select separate, optimized stimulus presentation orderings and timings for each participant. This procedure used AFNI `make_random_timing.py` to generate thousands of possible stimulus timings, and AFNI `3dDeconvolve` to select the timings that best supported deconvolving unique patterns of brain activity for each stimulus. Stimuli were presented using PsychoPy version 1.84.2 (Peirce, 2007). Participants were instructed to attend to the emotion content of the stimuli. During randomly interspersed catch trials (10 per run), participants used a button box to rate on a four-point scale whether the most recently presented stimulus had emotion content that was “more mixed” or “more pure.” To ensure familiarity with the stimuli, all fMRI participants had previously completed the emotion evaluation task.

#### fMRI acquisition

Participants were scanned at the Dartmouth Brain Imaging Center using a 3T Phillips Achieva Intera scanner with a 32-channel head coil. Functional images were acquired using an echo-planar sequence (35ms TE, 3000ms TR; 90° flip angle; 3×3×3mm resolution) with 192 dynamic scans per run. A high resolution T1-weighted anatomical scan (3.7 ms TE; 8200ms TR; .938x.938×1mm resolution) was acquired at the end of each scanning session. Sound was delivered using an over-ear headphone system. Foam padding was placed around participants’ heads to minimize motion.

#### fMRI preprocessing

Anatomical images were skull-stripped and aligned to the last TR of the last EPI image using AFNI `align_epi_anat.py`. EPI images were motion corrected and aligned to the last TR of the last EPI image using AFNI `3dvolreg`. Rigid body transformations for aligning participants’ anatomical and EPI images to the AFNI version of the MNI 152 ICBM template were calculated using AFNI @auto_tlrc. Alignment transformations were concatenated and applied in a single step using AFNI `3dAllineate`. EPI images were scaled to show percent signal change and concatenated. EPI images were not smoothed. TRs where inter-TR motion exceeded a euclidean norm threshold of .3 were censored, along with the immediately preceding TR. The general linear model was used to estimate BOLD-responses evoked by each of the 76 emotional stimulus classes using AFNI `3dREML-fit`. All six demeaned motion parameters as well as polynomial trends were included as regressors of no interest.

### Quantification and statistical analysis

#### Post hoc power analysis

Because the present study is the first to use the reported paradigm, we did not conduct an *a priori*/prospective power analysis. Because accurate assessment of effect size is impossible without stable patterns, we prioritized having a large number of fMRI trials per participant. The number of trials per stimulus class per participant was determined by consulting studies that used similar analysis methods (MVPA/RSA). E.g., Peelen et al. (2010) used 12 trials per class per participant for 18 participants, Kim et al. (2017) used 10 trials per class per participant for 20 participants, and the present study used 18 trials per class per participant for 20 participants. A *post hoc*/retrospective power analysis using G*Power 3.1 (Faul, Erdfelder, Lang, & Buchner, 2007) showed that this provided 85% power for music trials and 99% power for animation trials for the effects reported in Figure 3.

#### Representational similarity analysis

Representational similarity analysis (RSA) (Kriegeskorte et al., 2006, 2008) was conducted using PyMVPA (Hanke et al., 2009), Scikit-Learn (Pedregosa et al., 2012), NumPy (Oliphant, 2006), and SciPy (Jones, Oliphant, & Peterson, 2001). Stimulus feature representational distance matrices (RDMs) for each parameter (speed, irregularity/jitter, consonance/spikiness, ratio of big-to-small movements, ratio of upward-to-downward movements) were created by aggregating the Euclidean distances between the mean slider bar settings for each pair of emotions, including mixed emotions. Both music and animation stimuli were created using the same slider bar settings for each emotion, making it unnecessary to create modality-specific feature RDMs. Emotion RDMs were created by aggregating the Euclidean distances between the mean of each emotion judgment parameter in experiment 1 (Angry, Happy, Peaceful, Sad, and Scared) for each pair of emotions, including mixed emotions. Emotion judgments were averaged across music and animation, making it unnecessary to create modality-specific emotion judgment RDMs. Intermodal RDMs were built by calculating the full multi-modality RDM including both music and animation stimuli and selecting its lower-left square region (Figure 6B).

Representational similarity analysis was conducted separately for music trials, animation trials, and (for the intermodal analysis) music and animation trials together. Each analysis used a spherical searchlight with a 3-voxel (9mm) radius. In each searchlight sphere, music and animation neural RDMs were created by aggregating the ranked correlation distances (1-Spearman’s ρ) between the estimated stimulus-evoked pattern of BOLD activation for each emotion. The use of correlation distance ensured that the analysis would not mistake differences in the mean level of BOLD activity across music and animation trials for differences in representational similarity. Intermodal neural RDMs were created as described above, using neural data instead of stimulus features or emotion judgments (Figure 6B). The fit of the model to stimulus-evoked patterns of BOLD activation was assessed using multiple regression, with the ranked model RDMs as predictors and the neural RDM as the target. This produced coefficient of determination (*R*^2^) and β weight maps for each participant and each analysis. *R*^2^ values were adjusted using a permutation approach (similar to that of Peres-Neto, Legendre, Dray, & Borcard, 2006): Multiple regression was performed an additional 1000 times with randomly selected permutations of each predictor, and the mean *R*^2^ from this null distribution was subtracted from the reported *R*^2^ values 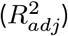. Multiple regression β weights reflect only the unique contribution of each predictor, resulting in β weight maps that do not reflect the shared contributions of correlated predictors. To assess the contribution of individual predictors we calculated the ranked correlation (Spearman’s ρ) of each predictor to the neural RDM.

All group-level statistics (including 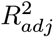, β weights, Spearman’s ρ, p-values, and any other values reported unless otherwise noted) were corrected for multiple comparisons using a maximum cluster mass sign-flipping permutation test performed with FSL randomise (Jenkinson, Beckmann, Behrens, Woolrich, & Smith, 2012; Nichols & Holmes, 2001), with a cluster-determining threshold of p=.01 and a family-wise error rate of .05.

#### Intermodal RSA

Intermodal RSA differed from the RSA analysis described above in that both the neural target RDM and the predictor RDMs used only between-modality distances, corresponding to the lower-left square region of the larger triangular RDM created using stimuli from both modalities (Figure 6B). If activity in a brain area was unrelated to stimuli presented in its non-preferred modality, then the intermodal neural RDM should be uncorrelated with the intermodal model RDMs. However, if a brain area was even weakly representing emotion content across modalities, then the intermodal neural RDM should be correlated with the intermodal model RDMs. Note that because this analysis only considered between-modality distances, it could not in principle have identified any modality-specific activity.

#### Model-free similarity analysis

To rule out the possibility that the identified brain regions were a good fit for the stimulus features and emotion judgments in the reported model, but did not truly share a representational geometry (i.e., were not directly similar to each other), we performed a permutation test of inter-region representational similarity. This test assessed whether the representations at the locations of peak model fit were more similar than representations at randomly selected locations. Analogously, the claim “San Francisco and Oakland are close to each other,” is weaker than the claim “San Francisco and Oakland are closer to each other than 95% of all pairs of American cities.” To build a null distribution, we randomly selected 2000 pairs of coordinates in the right hemisphere of the brain. For each coordinate pair, we measured the ranked correlation (Spearman’s ρ) of the mean neural RDM for music trials at the first coordinate and the mean neural RDM for animation trials at the second coordinate. The mean inter-region similarity in the null distribution was ρ=−.007, whereas the inter-region similarity at the locations of peak model fit was ρ=.68 (p<.001), more similar than any pair of coordinates in the null distribution.

#### Noise ceiling

The upper and lower bounds of the noise ceiling were calculated using an approach based on Nili et al. (2014), but adapted for use with multiple regression. The approach described by Nili et al. (2014) depends on a simple principle: for any dataset, the model that accounts for the most variance in the data will always be derived from the data itself. For correlation, this best-fitting model is the mean of the data. Given measurement error and individual differences across the dataset, no model could possibly outperform the mean, and so the model fit of the mean establishes a ceiling against which other models can be usefully compared. Analogously, for a multiple regression model with n predictors, the best-fitting model is the mean of the data along with the top n predictors identified using principal component analysis (PCA). No multiple regression model with the same number of predictors could possibly outperform this mean-and-PCA model. In the present study, the upper bound of the noise ceiling was calculated at each searchlight center by performing a multiple regression analysis that used the mean neural RDM and the top 10 principal components of the neural RDM as predictors. The lower bound of the noise ceiling was calculated using a leave-one-subject-out cross-validation approach: For each subject, the same multiple regression procedure was applied, but the mean neural RDM and the top 10 principal components were calculated with that subject left out.

#### Overlap maps

Overlap maps were created for each participant by identifying voxels where both music and animation model fits were significant at the individual level (permutation p<.05, uncorrected). Overlap maps were set to 1 if both model fits were significant, and 0 otherwise. Multiple comparisons correction of the overlap maps was performed at the group level (CDT=.01; FWER p=.05; see below), testing the proportion of individuals that showed overlap in a region against the null hypothesis that no participants showed overlap in that region.

#### Multiple comparisons correction

Group level maps were calculated and corrected for multiple comparisons using a maximum cluster mass sign-flipping permutation test FSL randomise (Jenkinson et al., 2012; Nichols & Holmes, 2001) (cluster-determining threshold p=.01; family-wise error rate p=.05). Tests for 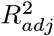 were 1-sided. Tests for β weights and Spearman’s ρ were two-sided. Maps were visualized using Nilearn (Abraham et al., 2014) and AFNI SUMA (Saad, Reynolds, Argall, Japee, & Cox, 2004).

#### Emotion judgments permutation procedure

For each emotion, we averaged participants’ emotion judgment ratings, yielding a class mean. We then calculated the Euclidean distance of each individual judgment to this mean, scaled by the maximum possible distance (determined by the limits of each slider), yielding a distribution of scaled distances to the mean for each stimulus class. A null distribution of scaled distances to the class means was created by applying this procedure 2000 times, each with a different permutation of the emotion labels over the whole dataset. Welch’s independent samples *t*-test was applied to test whether the observed distributions of scaled distances to class means differed from the null. This approach was chosen because it accounts for the simultaneous use of five rating scales and conservatively respects the dependency structure of the data.

