## Supplementary material for "Visual and auditory brain areas share a representational structure that supports emotion perception"

### Supplementary figures

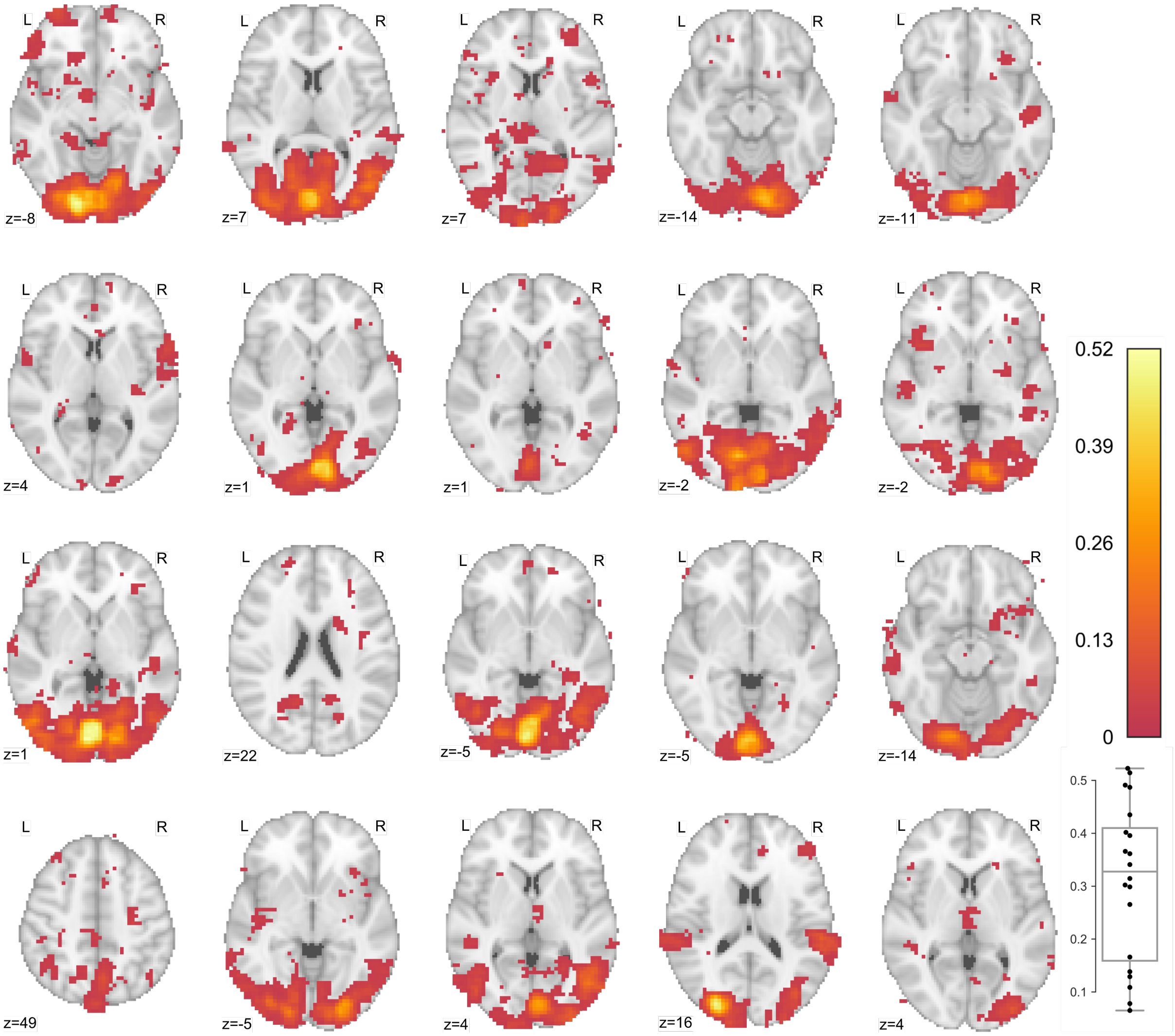


**Figure S1:** Peak model fit for animation trials for all 20 participants, related to Figure 5A. Brain maps show location of peak model fit in the axial plane, thresholded at p<.05 uncorrected. Box plot shows distribution of peak model fits across participants (mean $R^2_{adj}$=.31; 95% CI: .24–.38; t(19)=9.2; p<.001).

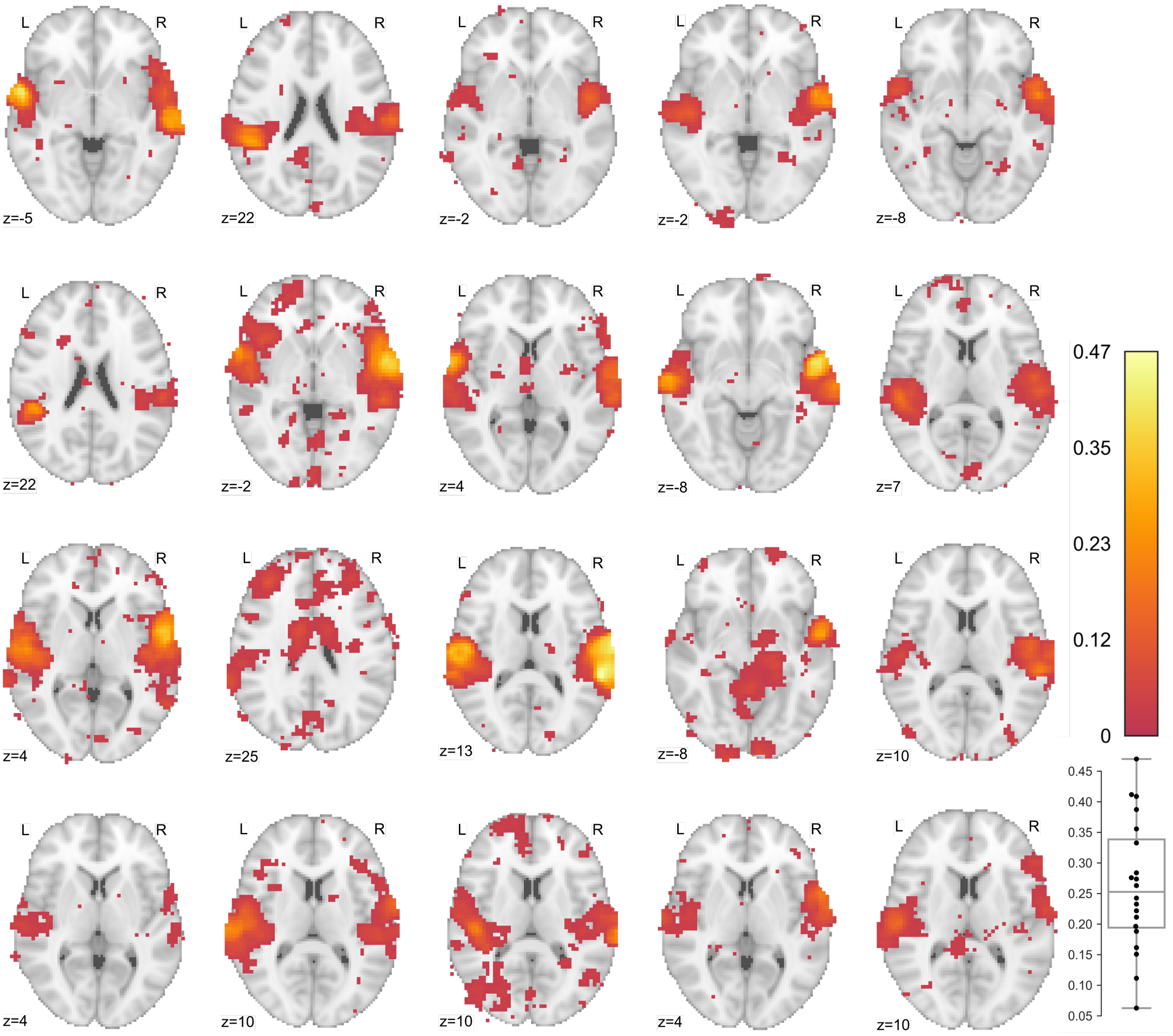


**Figure S2:** Peak model fit for music trials for all 20 participants, related to Figure 5B. Brain maps show location of peak model fit in the axial plane, thresholded at p<.05 uncorrected. Box plot shows distribution of peak model fits across participants (mean $R^2_{adj}$=.26; 95% CI: .21–.31; t(19)=10.95; p<.001).

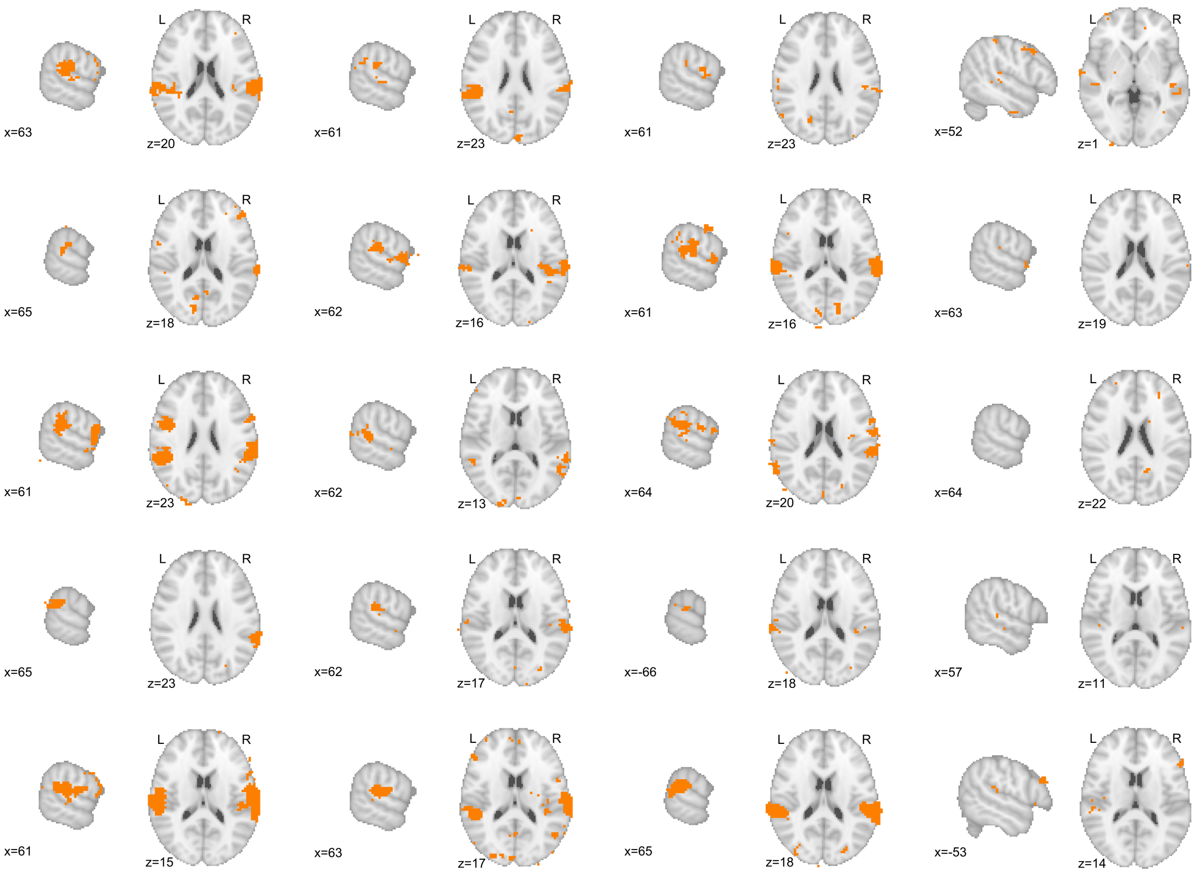


**Figure S3:** Supramodal representation maps for all 20 participants, related to Figure 6A. Masks show voxels with significant model fit ($p<.05$ uncorrected) during both animation and music trials. Cut points were selected to emphasize the presence or absence of overlap in bilateral posterior superior temporal gyrus.

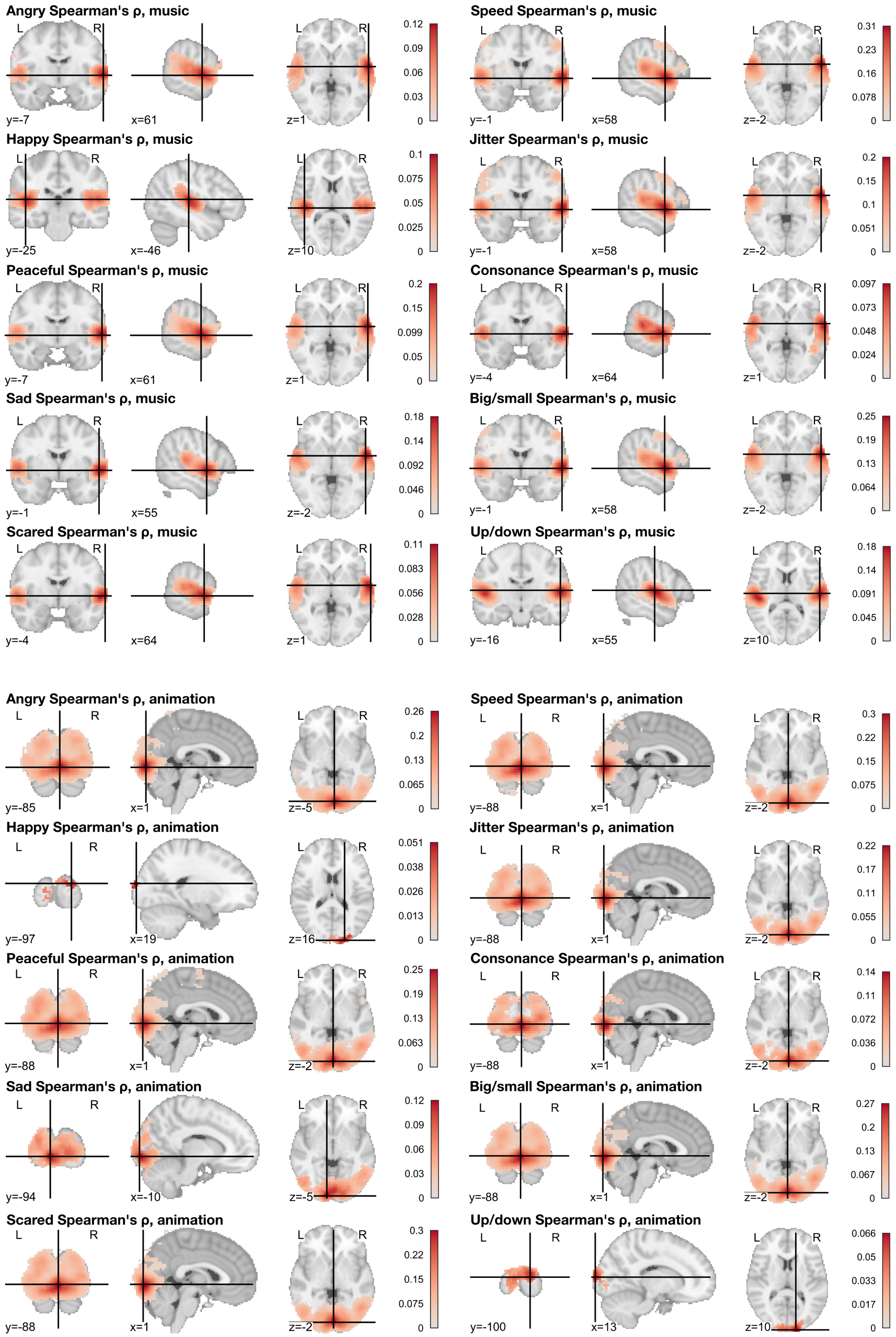


**Figure S4:** Parameter Spearman's ρ across participants, related to Figure 5. Top: model fit to music trials. Bottom: model fit to animation trials. Maps thresholded at voxelwise FWER=.05. Crosshairs indicate peak Spearman's ρ weight.

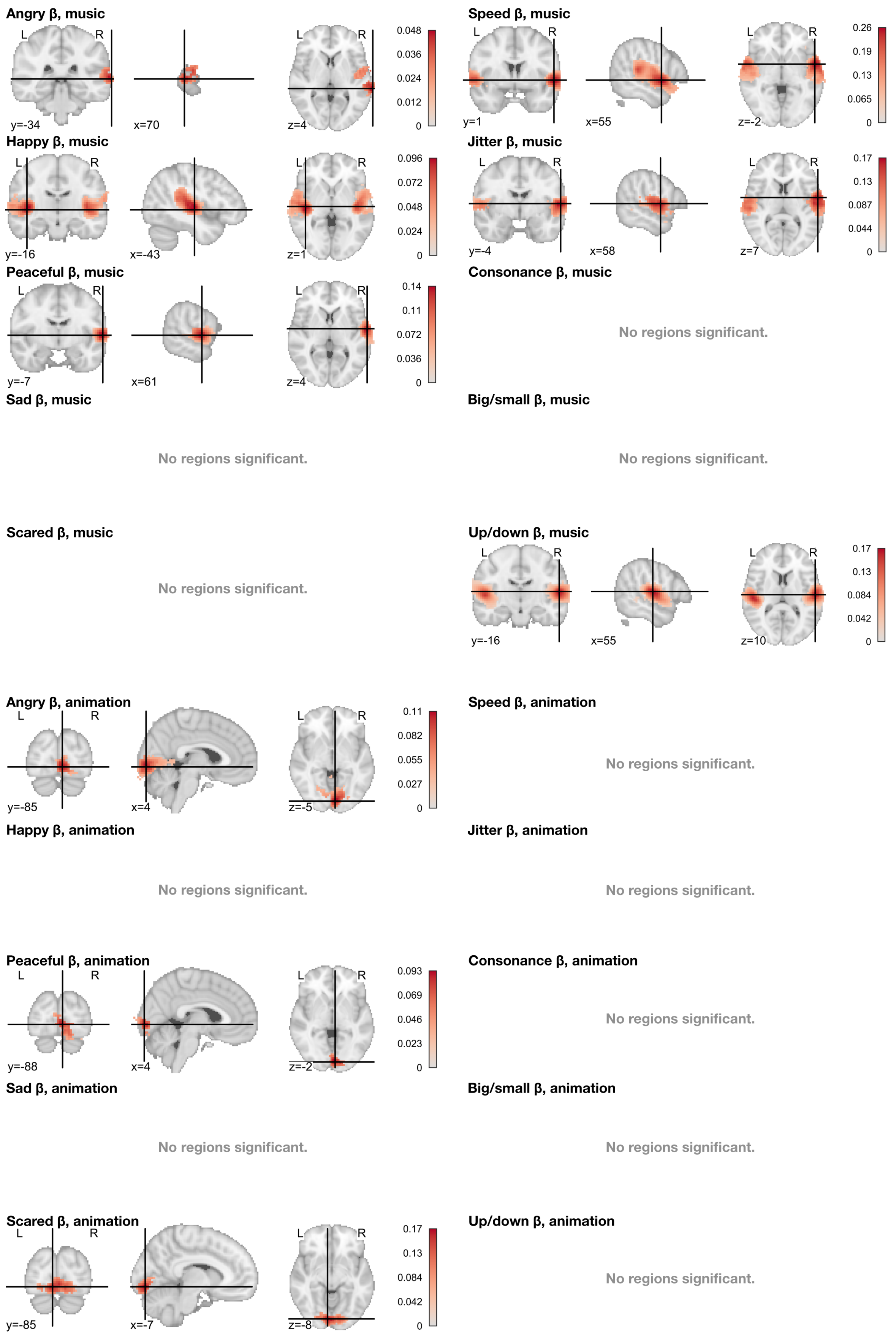


**Figure S5:** Parameter β weights across participants, related to Figure 5. Top: model fit to music trials. Bottom: model fit to animation trials. Maps thresholded at voxelwise FWER=.05. Crosshairs indicate peak β weight.

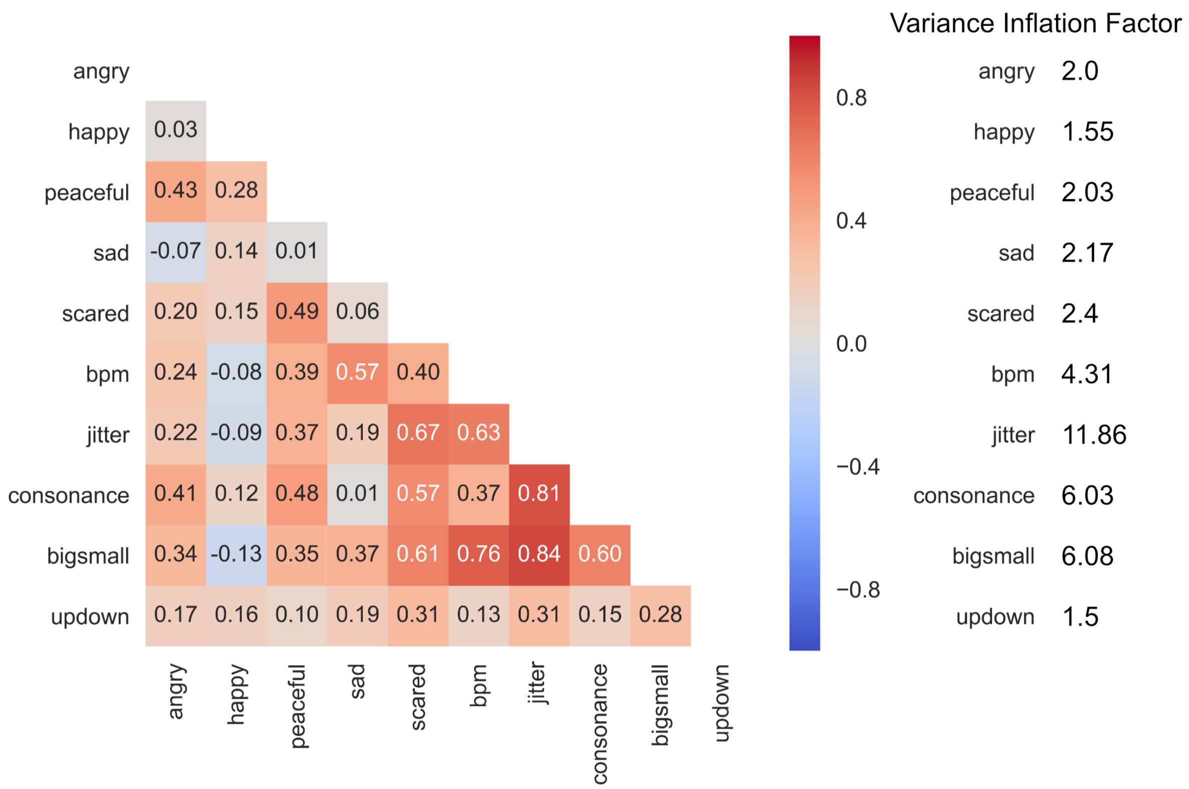


**Figure S6:** Model multicollinearity and variance inflation factors, related to Figure 5.

### Supplementary table

**Table S1:** Locations of peak model fits for individual predictors. Spearman's ρ reflects both the unique and shared contributions of the predictor, while β weights reflect only the unique contribution of the predictor.

| Parameter | x, y, z | Nearest atlas label[S1] | ρ | β | 95% CI | p |
|---|---|---|---|---|---|---|
| Anim. Angry | 2, -86, -5 | L Lingual gyrus, lingual part of the medial occipito-temporal gyrus, (O5) | .26 | | .18–.34 | < .001 |
| Anim. Angry | 50, -64, 1 | R Inferior temporal gyrus (T3) | .12 | | .09–.16 | < .001 |
| Anim. Angry | -22, -86, 28 | L Superior occipital gyrus (O1) | .11 | | .07–.16 | < .001 |
| Anim. Angry | 26, -82, 31 | R Superior occipital gyrus (O1) | .11 | | .07–.15 | < .001 |
| Anim. Angry | 64, -32, 25 | R Supramarginal gyrus | .10 | | .06–.13 | < .001 |
| Anim. Angry | -56, -38, 22 | L Planum temporale or temporal plane of the superior temporal gyrus | .08 | | .04–.12 | < .001 |
| Anim. Angry | -34, -50, 58 | L Superior parietal lobule (lateral part of P1) | .06 | | .03–.09 | < .001 |
| Anim. Angry | 34, -50, 58 | R Superior parietal lobule (lateral part of P1) | .06 | | .03–.08 | < .001 |

| Parameter | x, y, z | Nearest atlas label[S1] | ρ | β | 95% CI | p |
|---|---|---|---|---|---|---|
| Anim. Angry | 14, -28, 46 | R Marginal branch (or part) of the cingulate sulcus | .05 | | .02–.09 | < .001 |
| Anim. Happy | 20, -98, 16 | R Occipital pole | .05 | | .02–.08 | .013 |
| Anim. Happy | 38, -82, -14 | R Inferior occipital gyrus (O3) and sulcus | .04 | | .02–.06 | .013 |
| Anim. Happy | -20, -92, -8 | L Occipital pole | .03 | | .02–.05 | .013 |
| Anim. Peaceful | 2, -88, -2 | L Lingual gyrus, lingual part of the medial occipito-temporal gyrus, (O5) | .25 | | .18–.33 | < .001 |
| Anim. Peaceful | 50, -64, 1 | R Inferior temporal gyrus (T3) | .14 | | .10–.19 | < .001 |
| Anim. Peaceful | -46, -70, 4 | L Middle occipital gyrus (O2, lateral occipital gyrus) | .14 | | .09–.19 | < .001 |
| Anim. Peaceful | 64, -32, 25 | R Supramarginal gyrus | .12 | | .07–.17 | < .001 |
| Anim. Peaceful | -52, -38, 22 | L Planum temporale or temporal plane of the superior temporal gyrus | .09 | | .04–.14 | < .001 |
| Anim. Peaceful | -14, -26, 40 | L Marginal branch (or part) of the cingulate sulcus | .07 | | .03–.11 | .042 |
| Anim. Peaceful | 38, -52, 61 | R Superior parietal lobule (lateral part of P1) | .07 | | .04–.09 | < .001 |
| Anim. Peaceful | 52, 8, 34 | R Precentral gyrus | .06 | | .03–.10 | < .001 |
| Anim. Peaceful | 10, -28, 46 | R Marginal branch (or part) of the cingulate sulcus | .06 | | .02–.10 | < .001 |
| Anim. Peaceful | -34, -56, 61 | L Superior parietal lobule (lateral part of P1) | .06 | | .03–.08 | < .001 |
| Anim. Peaceful | -28, -70, -53 | L Lateral occipito-temporal gyrus (fusiform gyrus, O4-T4) | .05 | | .03–.08 | < .001 |
| Anim. Peaceful | 8, -10, 70 | R Superior frontal gyrus (F1) | .05 | | .02–.08 | .042 |
| Anim. Peaceful | 44, 40, 19 | R Middle frontal gyrus (F2) | .04 | | .02–.06 | .047 |
| Anim. Peaceful | -58, 8, 37 | L Precentral gyrus | .04 | | .02–.06 | < .001 |
| Anim. Sad | -10, -94, -5 | L Occipital pole | .12 | | .07–.17 | < .001 |
| Anim. Sad | 46, -68, 1 | R Inferior occipital gyrus (O3) and sulcus | .07 | | .04–.10 | < .001 |
| Anim. Sad | 64, -28, 19 | R Planum temporale or temporal plane of the superior temporal gyrus | .07 | | .04–.09 | < .001 |
| Anim. Sad | -58, -32, 19 | L Supramarginal gyrus | .04 | | .02–.07 | .018 |

| Parameter | x, y, z | Nearest atlas label[S1] | ρ | β | 95% CI | p |
|---|---|---|---|---|---|---|
| Anim. Sad | 52, 8, 34 | R Precentral gyrus | .04 | | .01–.06 | < .001 |
| Anim. Scared | 2, -88, -2 | L Lingual gyrus, lingual part of the medial occipito-temporal gyrus, (O5) | .30 | | .21–.39 | < .001 |
| Anim. Scared | 50, -64, 1 | R Inferior temporal gyrus (T3) | .15 | | .11–.20 | < .001 |
| Anim. Scared | -22, -86, 28 | L Superior occipital gyrus (O1) | .14 | | .08–.20 | < .001 |
| Anim. Scared | 22, -82, 31 | R Superior occipital gyrus (O1) | .14 | | .09–.19 | < .001 |
| Anim. Scared | 62, -34, 25 | R Planum temporale or temporal plane of the superior temporal gyrus | .13 | | .08–.17 | < .001 |
| Anim. Scared | -52, -38, 22 | L Planum temporale or temporal plane of the superior temporal gyrus | .11 | | .05–.16 | < .001 |
| Anim. Scared | -34, -56, 61 | L Superior parietal lobule (lateral part of P1) | .08 | | .04–.11 | < .001 |
| Anim. Scared | -58, 10, 37 | L Precentral gyrus | .04 | | .01–.07 | < .001 |
| Anim. Speed | 2, -88, -2 | L Lingual gyrus, lingual part of the medial occipito-temporal gyrus, (O5) | .30 | | .21–.39 | < .001 |
| Anim. Speed | 50, -64, 1 | R Inferior temporal gyrus (T3) | .17 | | .12–.22 | < .001 |
| Anim. Speed | -22, -86, 28 | L Superior occipital gyrus (O1) | .15 | | .09–.21 | < .001 |
| Anim. Speed | 26, -80, 28 | R Superior occipital gyrus (O1) | .14 | | .10–.19 | < .001 |
| Anim. Speed | 64, -32, 22 | R Supramarginal gyrus | .14 | | .08–.19 | < .001 |
| Anim. Speed | -52, -38, 22 | L Planum temporale or temporal plane of the superior temporal gyrus | .11 | | .05–.16 | < .001 |
| Anim. Speed | 34, -52, 58 | R Superior parietal lobule (lateral part of P1) | .07 | | .04–.10 | < .001 |
| Anim. Speed | -34, -56, 61 | L Superior parietal lobule (lateral part of P1) | .07 | | .04–.10 | < .001 |
| Anim. Speed | 52, 8, 34 | R Precentral gyrus | .06 | | .03–.10 | < .001 |
| Anim. Speed | -28, -70, -53 | L Lateral occipito-temporal gyrus (fusiform gyrus, O4-T4) | .06 | | .03–.09 | < .001 |
| Anim. Speed | -56, 10, 10 | L Opercular part of the inferior frontal gyrus | .04 | | .02–.07 | < .001 |
| Anim. Jitter | 2, -88, -2 | L Lingual gyrus, lingual part of the medial occipito-temporal gyrus, (O5) | .22 | | .16–.28 | < .001 |

| Parameter | x, y, z | Nearest atlas label[S1] | ρ | β | 95% CI | p |
|---|---|---|---|---|---|---|
| Anim. Jitter | 50, -64, 1 | R Inferior temporal gyrus (T3) | .11 | | .08–.14 | < .001 |
| Anim. Jitter | -50, -70, 1 | L Inferior temporal gyrus (T3) | .11 | | .07–.15 | < .001 |
| Anim. Jitter | 62, -32, 25 | R Planum temporale or temporal plane of the superior temporal gyrus | .09 | | .05–.13 | < .001 |
| Anim. Jitter | -56, -40, 22 | L Planum temporale or temporal plane of the superior temporal gyrus | .06 | | .03–.09 | < .001 |
| Anim. Jitter | 38, -52, 58 | R Superior parietal lobule (lateral part of P1) | .06 | | .03–.08 | < .001 |
| Anim. Consonance | 2, -88, -2 | L Lingual gyrus, lingual part of the medial occipito-temporal gyrus, (O5) | .14 | | .10–.19 | < .001 |
| Anim. Consonance | -52, -74, 1 | L Middle occipital gyrus (O2, lateral occipital gyrus) | .08 | | .05–.11 | < .001 |
| Anim. Consonance | -28, -82, 31 | L Middle occipital gyrus (O2, lateral occipital gyrus) | .06 | | .03–.09 | < .001 |
| Anim. Consonance | 62, -32, 25 | R Planum temporale or temporal plane of the superior temporal gyrus | .05 | | .02–.08 | < .001 |
| Anim. Consonance | -62, -40, 25 | L Planum temporale or temporal plane of the superior temporal gyrus | .04 | | .02–.06 | < .001 |
| Anim. Big/small | 2, -88, -2 | L Lingual gyrus, lingual part of the medial occipito-temporal gyrus, (O5) | .27 | | .19–.35 | < .001 |
| Anim. Big/small | 50, -64, 1 | R Inferior temporal gyrus (T3) | .14 | | .10–.18 | < .001 |
| Anim. Big/small | -22, -86, 28 | L Superior occipital gyrus (O1) | .12 | | .06–.17 | < .001 |
| Anim. Big/small | 28, -82, 31 | R Superior occipital gyrus (O1) | .12 | | .08–.16 | < .001 |
| Anim. Big/small | 62, -32, 25 | R Planum temporale or temporal plane of the superior temporal gyrus | .10 | | .06–.14 | < .001 |
| Anim. Big/small | -56, -34, 25 | L Supramarginal gyrus | .07 | | .03–.11 | < .001 |
| Anim. Big/small | 32, -52, 58 | R Superior parietal lobule (lateral part of P1) | .06 | | .03–.09 | < .001 |
| Anim. Big/small | 52, 8, 34 | R Precentral gyrus | .06 | | .02–.09 | < .001 |
| Anim. Big/small | -34, -56, 64 | L Superior parietal lobule (lateral part of P1) | .05 | | .03–.08 | < .001 |
| Anim. Up/down | 14, -100, 10 | R Occipital pole | .07 | | .03–.10 | .004 |
| Mus. Angry | 62, -8, 1 | R Lateral aspect of the superior temporal gyrus | .12 | | .10–.14 | < .001 |

| Parameter | x, y, z | Nearest atlas label[S1] | ρ | β | 95% CI | p |
|---|---|---|---|---|---|---|
| Mus. Angry | -56, -14, 7 | L Anterior transverse temporal gyrus (of Heschl) | .08 | | .04–.11 | < .001 |
| Mus. Angry | 2, -64, 55 | R Precuneus (medial part of P1) | .03 | | .01–.05 | .016 |
| Mus. Angry | -10, -68, 25 | L Precuneus (medial part of P1) | .03 | | .01–.05 | .016 |
| Mus. Happy | -46, -26, 10 | L Anterior transverse temporal gyrus (of Heschl) | .10 | | .07–.13 | < .001 |
| Mus. Happy | 46, -22, 10 | R Anterior transverse temporal gyrus (of Heschl) | .08 | | .06–.10 | < .001 |
| Mus. Peaceful | 62, -8, 1 | R Lateral aspect of the superior temporal gyrus | .20 | | .16–.24 | < .001 |
| Mus. Peaceful | -62, -20, 10 | L Planum temporale or temporal plane of the superior temporal gyrus | .13 | | .09–.18 | < .001 |
| Mus. Sad | 56, -2, -2 | R Planum polare of the superior temporal gyrus | .18 | | .13–.24 | < .001 |
| Mus. Sad | -58, -2, 1 | L Lateral aspect of the superior temporal gyrus | .13 | | .09–.17 | < .001 |
| Mus. Scared | 64, -4, 1 | R Lateral aspect of the superior temporal gyrus | .11 | | .08–.14 | < .001 |
| Mus. Scared | -58, -4, 1 | L Lateral aspect of the superior temporal gyrus | .08 | | .05–.11 | < .001 |
| Mus. Speed | 58, -2, -2 | R Lateral aspect of the superior temporal gyrus | .31 | | .24–.38 | < .001 |
| Mus. Speed | 64, -32, 13 | R Lateral aspect of the superior temporal gyrus | .22 | | .15–.30 | < .001 |
| Mus. Speed | -68, -16, 7 | L Lateral aspect of the superior temporal gyrus | .22 | | .15–.28 | < .001 |
| Mus. Speed | 52, -2, 46 | R Precentral gyrus | .10 | | .05–.14 | < .001 |
| Mus. Speed | -56, -4, 49 | L Precentral gyrus | .08 | | .04–.12 | < .001 |
| Mus. Speed | -16, -70, 28 | L Parieto-occipital sulcus (or fissure) | .05 | | .03–.07 | .011 |
| Mus. Speed | -46, -86, 19 | L Middle occipital gyrus (O2, lateral occipital gyrus) | .04 | | .02–.06 | .011 |
| Mus. Jitter | 58, -2, -2 | R Lateral aspect of the superior temporal gyrus | .20 | | .16–.25 | < .001 |
| Mus. Jitter | -62, -16, 4 | L Lateral aspect of the superior temporal gyrus | .15 | | .11–.20 | < .001 |
| Mus. Jitter | 52, 2, 46 | R Precentral gyrus | .08 | | .05–.11 | < .001 |
| Mus. Jitter | -56, -8, 52 | L Precentral gyrus | .06 | | .03–.08 | < .001 |

| Parameter | x, y, z | Nearest atlas label[S1] | ρ | β | 95% CI | p |
|---|---|---|---|---|---|---|
| Mus. Jitter | -16, -68, 28 | L Parieto-occipital sulcus (or fissure) | .05 | | .03–.07 | .012 |
| Mus. Jitter | 26, -80, 31 | R Superior occipital gyrus (O1) | .03 | | .01–.05 | .012 |
| Mus. Consonance | 64, -4, 1 | R Lateral aspect of the superior temporal gyrus | .10 | | .07–.13 | < .001 |
| Mus. Consonance | -62, -16, 7 | L Lateral aspect of the superior temporal gyrus | .08 | | .04–.11 | < .001 |
| Mus. Consonance | -56, -38, 16 | L Planum temporale or temporal plane of the superior temporal gyrus | .07 | | .04–.09 | < .001 |
| Mus. Big/small | 58, -2, -2 | R Lateral aspect of the superior temporal gyrus | .25 | | .20–.31 | < .001 |
| Mus. Big/small | -64, -16, 7 | L Lateral aspect of the superior temporal gyrus | .18 | | .13–.24 | < .001 |
| Mus. Big/small | 52, -2, 46 | R Precentral gyrus | .09 | | .05–.13 | < .001 |
| Mus. Big/small | -56, -4, 49 | L Precentral gyrus | .06 | | .03–.09 | < .001 |
| Mus. Big/small | -16, -68, 28 | L Parieto-occipital sulcus (or fissure) | .04 | | .03–.06 | .012 |
| Mus. Up/down | 56, -16, 10 | R Anterior transverse temporal gyrus (of Heschl) | .18 | | .14–.22 | < .001 |
| Mus. Up/down | -50, -26, 10 | L Anterior transverse temporal gyrus (of Heschl) | .17 | | .14–.21 | < .001 |
| Anim. Happy | 20, -98, 16 | R Occipital pole | .05 | | .02–.08 | .013 |
| Anim. Happy | 38, -82, -14 | R Inferior occipital gyrus (O3) and sulcus | .04 | | .02–.06 | .013 |
| Anim. Happy | -20, -92, -8 | L Occipital pole | .03 | | .02–.05 | .013 |
| Anim. Peaceful | 2, -88, -2 | L Lingual gyrus, lingual part of the medial occipito-temporal gyrus, (O5) | .25 | | .18–.33 | < .001 |
| Anim. Peaceful | 50, -64, 1 | R Inferior temporal gyrus (T3) | .14 | | .10–.19 | < .001 |
| Anim. Peaceful | -46, -70, 4 | L Middle occipital gyrus (O2, lateral occipital gyrus) | .14 | | .09–.19 | < .001 |
| Anim. Peaceful | 64, -32, 25 | R Supramarginal gyrus | .12 | | .07–.17 | < .001 |
| Anim. Peaceful | -52, -38, 22 | L Planum temporale or temporal plane of the superior temporal gyrus | .09 | | .04–.14 | < .001 |
| Anim. Peaceful | -14, -26, 40 | L Marginal branch (or part) of the cingulate sulcus | .07 | | .03–.11 | .042 |
| Anim. Peaceful | 38, -52, 61 | R Superior parietal lobule (lateral part of P1) | .07 | | .04–.09 | < .001 |

| Parameter | x, y, z | Nearest atlas label[S1] | ρ | β | 95% CI | p |
|---|---|---|---|---|---|---|
| Anim. Peaceful | 52, 8, 34 | R Precentral gyrus | .06 | | .03–.10 | < .001 |
| Anim. Peaceful | 10, -28, 46 | R Marginal branch (or part) of the cingulate sulcus | .06 | | .02–.10 | < .001 |
| Anim. Peaceful | -34, -56, 61 | L Superior parietal lobule (lateral part of P1) | .06 | | .03–.08 | < .001 |
| Anim. Peaceful | -28, -70, -53 | L Lateral occipito-temporal gyrus (fusiform gyrus, O4-T4) | .05 | | .03–.08 | < .001 |
| Anim. Peaceful | 8, -10, 70 | R Superior frontal gyrus (F1) | .05 | | .02–.08 | .042 |
| Anim. Peaceful | 44, 40, 19 | R Middle frontal gyrus (F2) | .04 | | .02–.06 | .047 |
| Anim. Peaceful | -58, 8, 37 | L Precentral gyrus | .04 | | .02–.06 | < .001 |
| Anim. Sad | -10, -94, -5 | L Occipital pole | .12 | | .07–.17 | < .001 |
| Anim. Sad | 46, -68, 1 | R Inferior occipital gyrus (O3) and sulcus | .07 | | .04–.10 | < .001 |
| Anim. Sad | 64, -28, 19 | R Planum temporale or temporal plane of the superior temporal gyrus | .07 | | .04–.09 | < .001 |
| Anim. Sad | -58, -32, 19 | L Supramarginal gyrus | .04 | | .02–.07 | .018 |
| Anim. Sad | 52, 8, 34 | R Precentral gyrus | .04 | | .01–.06 | < .001 |
| Anim. Scared | 2, -88, -2 | L Lingual gyrus, lingual part of the medial occipito-temporal gyrus, (O5) | .30 | | .21–.39 | < .001 |
| Anim. Scared | 50, -64, 1 | R Inferior temporal gyrus (T3) | .15 | | .11–.20 | < .001 |
| Anim. Scared | -22, -86, 28 | L Superior occipital gyrus (O1) | .14 | | .08–.20 | < .001 |
| Anim. Scared | 22, -82, 31 | R Superior occipital gyrus (O1) | .14 | | .09–.19 | < .001 |
| Anim. Scared | 62, -34, 25 | R Planum temporale or temporal plane of the superior temporal gyrus | .13 | | .08–.17 | < .001 |
| Anim. Scared | -52, -38, 22 | L Planum temporale or temporal plane of the superior temporal gyrus | .11 | | .05–.16 | < .001 |
| Anim. Scared | -34, -56, 61 | L Superior parietal lobule (lateral part of P1) | .08 | | .04–.11 | < .001 |
| Anim. Scared | -58, 10, 37 | L Precentral gyrus | .04 | | .01–.07 | < .001 |
| Anim. Speed | 2, -88, -2 | L Lingual gyrus, lingual part of the medial occipito-temporal gyrus, (O5) | .30 | | .21–.39 | < .001 |
| Anim. Speed | 50, -64, 1 | R Inferior temporal gyrus (T3) | .17 | | .12–.22 | < .001 |

| Parameter | x, y, z | Nearest atlas label[S1] | ρ | β | 95% CI | p |
|---|---|---|---|---|---|---|
| Anim. Speed | -22, -86, 28 | L Superior occipital gyrus (O1) | .15 | | .09–.21 | < .001 |
| Anim. Speed | 26, -80, 28 | R Superior occipital gyrus (O1) | .14 | | .10–.19 | < .001 |
| Anim. Speed | 64, -32, 22 | R Supramarginal gyrus | .14 | | .08–.19 | < .001 |
| Anim. Speed | -52, -38, 22 | L Planum temporale or temporal plane of the superior temporal gyrus | .11 | | .05–.16 | < .001 |
| Anim. Speed | 34, -52, 58 | R Superior parietal lobule (lateral part of P1) | .07 | | .04–.10 | < .001 |
| Anim. Speed | -34, -56, 61 | L Superior parietal lobule (lateral part of P1) | .07 | | .04–.10 | < .001 |
| Anim. Speed | 52, 8, 34 | R Precentral gyrus | .06 | | .03–.10 | < .001 |
| Anim. Speed | -28, -70, -53 | L Lateral occipito-temporal gyrus (fusiform gyrus, O4-T4) | .06 | | .03–.09 | < .001 |
| Anim. Speed | -56, 10, 10 | L Opercular part of the inferior frontal gyrus | .04 | | .02–.07 | < .001 |
| Anim. Jitter | 2, -88, -2 | L Lingual gyrus, lingual part of the medial occipito-temporal gyrus, (O5) | .22 | | .16–.28 | < .001 |
| Anim. Jitter | 50, -64, 1 | R Inferior temporal gyrus (T3) | .11 | | .08–.14 | < .001 |
| Anim. Jitter | -50, -70, 1 | L Inferior temporal gyrus (T3) | .11 | | .07–.15 | < .001 |
| Anim. Jitter | 62, -32, 25 | R Planum temporale or temporal plane of the superior temporal gyrus | .09 | | .05–.13 | < .001 |
| Anim. Jitter | -56, -40, 22 | L Planum temporale or temporal plane of the superior temporal gyrus | .06 | | .03–.09 | < .001 |
| Anim. Jitter | 38, -52, 58 | R Superior parietal lobule (lateral part of P1) | .06 | | .03–.08 | < .001 |
| Anim. Consonance | 2, -88, -2 | L Lingual gyrus, lingual part of the medial occipito-temporal gyrus, (O5) | .14 | | .10–.19 | < .001 |
| Anim. Consonance | -52, -74, 1 | L Middle occipital gyrus (O2, lateral occipital gyrus) | .08 | | .05–.11 | < .001 |
| Anim. Consonance | -28, -82, 31 | L Middle occipital gyrus (O2, lateral occipital gyrus) | .06 | | .03–.09 | < .001 |
| Anim. Consonance | 62, -32, 25 | R Planum temporale or temporal plane of the superior temporal gyrus | .05 | | .02–.08 | < .001 |
| Anim. Consonance | -62, -40, 25 | L Planum temporale or temporal plane of the superior temporal gyrus | .04 | | .02–.06 | < .001 |
| Anim. Big/small | 2, -88, -2 | L Lingual gyrus, lingual part of the medial occipito-temporal gyrus, (O5) | .27 | | .19–.35 | < .001 |

| Parameter | x, y, z | Nearest atlas label[S1] | ρ | β | 95% CI | p |
|---|---|---|---|---|---|---|
| Anim. Big/small | 50, -64, 1 | R Inferior temporal gyrus (T3) | .14 | | .10–.18 | < .001 |
| Anim. Big/small | -22, -86, 28 | L Superior occipital gyrus (O1) | .12 | | .06–.17 | < .001 |
| Anim. Big/small | 28, -82, 31 | R Superior occipital gyrus (O1) | .12 | | .08–.16 | < .001 |
| Anim. Big/small | 62, -32, 25 | R Planum temporale or temporal plane of the superior temporal gyrus | .10 | | .06–.14 | < .001 |
| Anim. Big/small | -56, -34, 25 | L Supramarginal gyrus | .07 | | .03–.11 | < .001 |
| Anim. Big/small | 32, -52, 58 | R Superior parietal lobule (lateral part of P1) | .06 | | .03–.09 | < .001 |
| Anim. Big/small | 52, 8, 34 | R Precentral gyrus | .06 | | .02–.09 | < .001 |
| Anim. Big/small | -34, -56, 64 | L Superior parietal lobule (lateral part of P1) | .05 | | .03–.08 | < .001 |
| Anim. Up/down | 14, -100, 10 | R Occipital pole | .07 | | .03–.10 | .004 |
| Mus. Angry | 62, -8, 1 | R Lateral aspect of the superior temporal gyrus | .12 | | .10–.14 | < .001 |
| Mus. Angry | -56, -14, 7 | L Anterior transverse temporal gyrus (of Heschl) | .08 | | .04–.11 | < .001 |
| Mus. Angry | 2, -64, 55 | R Precuneus (medial part of P1) | .03 | | .01–.05 | .016 |
| Mus. Angry | -10, -68, 25 | L Precuneus (medial part of P1) | .03 | | .01–.05 | .016 |
| Mus. Happy | -46, -26, 10 | L Anterior transverse temporal gyrus (of Heschl) | .10 | | .07–.13 | < .001 |
| Mus. Happy | 46, -22, 10 | R Anterior transverse temporal gyrus (of Heschl) | .08 | | .06–.10 | < .001 |
| Mus. Peaceful | 62, -8, 1 | R Lateral aspect of the superior temporal gyrus | .20 | | .16–.24 | < .001 |
| Mus. Peaceful | -62, -20, 10 | L Planum temporale or temporal plane of the superior temporal gyrus | .13 | | .09–.18 | < .001 |
| Mus. Sad | 56, -2, -2 | R Planum polare of the superior temporal gyrus | .18 | | .13–.24 | < .001 |
| Mus. Sad | -58, -2, 1 | L Lateral aspect of the superior temporal gyrus | .13 | | .09–.17 | < .001 |
| Mus. Scared | 64, -4, 1 | R Lateral aspect of the superior temporal gyrus | .11 | | .08–.14 | < .001 |
| Mus. Scared | -58, -4, 1 | L Lateral aspect of the superior temporal gyrus | .08 | | .05–.11 | < .001 |

| Parameter | x, y, z | Nearest atlas label[S1] | ρ | β | 95% CI | p |
|---|---|---|---|---|---|---|
| Mus. Speed | 58, -2, -2 | R Lateral aspect of the superior temporal gyrus | .31 | | .24–.38 | < .001 |
| Mus. Speed | 64, -32, 13 | R Lateral aspect of the superior temporal gyrus | .22 | | .15–.30 | < .001 |
| Mus. Speed | -68, -16, 7 | L Lateral aspect of the superior temporal gyrus | .22 | | .15–.28 | < .001 |
| Mus. Speed | 52, -2, 46 | R Precentral gyrus | .10 | | .05–.14 | < .001 |
| Mus. Speed | -56, -4, 49 | L Precentral gyrus | .08 | | .04–.12 | < .001 |
| Mus. Speed | -16, -70, 28 | L Parieto-occipital sulcus (or fissure) | .05 | | .03–.07 | .011 |
| Mus. Speed | -46, -86, 19 | L Middle occipital gyrus (O2, lateral occipital gyrus) | .04 | | .02–.06 | .011 |
| Mus. Jitter | 58, -2, -2 | R Lateral aspect of the superior temporal gyrus | .20 | | .16–.25 | < .001 |
| Mus. Jitter | -62, -16, 4 | L Lateral aspect of the superior temporal gyrus | .15 | | .11–.20 | < .001 |
| Mus. Jitter | 52, 2, 46 | R Precentral gyrus | .08 | | .05–.11 | < .001 |
| Mus. Jitter | -56, -8, 52 | L Precentral gyrus | .06 | | .03–.08 | < .001 |
| Mus. Jitter | -16, -68, 28 | L Parieto-occipital sulcus (or fissure) | .05 | | .03–.07 | .012 |
| Mus. Jitter | 26, -80, 31 | R Superior occipital gyrus (O1) | .03 | | .01–.05 | .012 |
| Mus. Consonance | 64, -4, 1 | R Lateral aspect of the superior temporal gyrus | .10 | | .07–.13 | < .001 |
| Mus. Consonance | -62, -16, 7 | L Lateral aspect of the superior temporal gyrus | .08 | | .04–.11 | < .001 |
| Mus. Consonance | -56, -38, 16 | L Planum temporale or temporal plane of the superior temporal gyrus | .07 | | .04–.09 | < .001 |
| Mus. Big/small | 58, -2, -2 | R Lateral aspect of the superior temporal gyrus | .25 | | .20–.31 | < .001 |
| Mus. Big/small | -64, -16, 7 | L Lateral aspect of the superior temporal gyrus | .18 | | .13–.24 | < .001 |
| Mus. Big/small | 52, -2, 46 | R Precentral gyrus | .09 | | .05–.13 | < .001 |
| Mus. Big/small | -56, -4, 49 | L Precentral gyrus | .06 | | .03–.09 | < .001 |
| Mus. Big/small | -16, -68, 28 | L Parieto-occipital sulcus (or fissure) | .04 | | .03–.06 | .012 |
| Mus. Up/down | 56, -16, 10 | R Anterior transverse temporal gyrus (of Heschl) | .18 | | .14–.22 | < .001 |

| Parameter | x, y, z | Nearest atlas label[S1] | ρ | β | 95% CI | p |
|---|---|---|---|---|---|---|
| Mus. Up/down | -50, -26, 10 | L Anterior transverse temporal gyrus (of Heschl) | .17 | | .14–.21 | < .001 |
| Anim. Angry | 4, -86, -5 | R Lingual gyrus, lingual part of the medial occipito-temporal gyrus, (O5) | | .11 | .07–.15 | .002 |
| Anim. Peaceful | 4, -88, -2 | R Lingual gyrus, lingual part of the medial occipito-temporal gyrus, (O5) | | .09 | .05–.14 | .009 |
| Anim. Scared | -8, -86, -8 | L Lingual gyrus, lingual part of the medial occipito-temporal gyrus, (O5) | | .17 | .09–.24 | < .001 |
| Mus. Angry | 70, -34, 4 | R Lateral aspect of the superior temporal gyrus | | .05 | .03–.06 | .012 |
| Mus. Angry | 58, 2, -2 | R Lateral aspect of the superior temporal gyrus | | .04 | .03–.06 | .005 |
| Mus. Happy | -44, -16, 1 | L Inferior segment of the circular sulcus of the insula | | .10 | .06–.13 | < .001 |
| Mus. Happy | 56, -28, 10 | R Planum temporale or temporal plane of the superior temporal gyrus | | .08 | .05–.11 | < .001 |
| Mus. Peaceful | 62, -8, 4 | R Lateral aspect of the superior temporal gyrus | | .14 | .10–.18 | < .001 |
| Mus. Speed | 56, 2, -2 | R Planum polare of the superior temporal gyrus | | .26 | .19–.33 | < .001 |
| Mus. Speed | 62, -32, 13 | R Planum temporale or temporal plane of the superior temporal gyrus | | .24 | .16–.32 | < .001 |
| Mus. Speed | -62, 4, 1 | L Lateral aspect of the superior temporal gyrus | | .21 | .12–.30 | < .001 |
| Mus. Speed | -58, -38, 16 | L Planum temporale or temporal plane of the superior temporal gyrus | | .18 | .12–.24 | < .001 |
| Mus. Jitter | 58, -4, 7 | R Opercular part of the inferior frontal gyrus | | .17 | .14–.21 | < .001 |
| Mus. Jitter | -62, -10, 1 | L Lateral aspect of the superior temporal gyrus | | .13 | .08–.18 | .002 |
| Mus. Up/down | 56, -16, 10 | R Anterior transverse temporal gyrus (of Heschl) | | .17 | .13–.21 | < .001 |
| Mus. Up/down | -52, -22, 10 | L Anterior transverse temporal gyrus (of Heschl) | | .16 | .12–.20 | < .001 |
| Anim. Peaceful | 4, -88, -2 | R Lingual gyrus, lingual part of the medial occipito-temporal gyrus, (O5) | | .09 | .05–.14 | .009 |
| Anim. Scared | -8, -86, -8 | L Lingual gyrus, lingual part of the medial occipito-temporal gyrus, (O5) | | .17 | .09–.24 | < .001 |
| Mus. Angry | 70, -34, 4 | R Lateral aspect of the superior temporal gyrus | | .05 | .03–.06 | .012 |
| Mus. Angry | 58, 2, -2 | R Lateral aspect of the superior temporal gyrus | | .04 | .03–.06 | .005 |

| Parameter | x, y, z | Nearest atlas label[S1] | ρ | β | 95% CI | p |
|---|---|---|---|---|---|---|
| Mus. Happy | -44, -16, 1 | L Inferior segment of the circular sulcus of the insula | | .10 | .06–.13 | < .001 |
| Mus. Happy | 56, -28, 10 | R Planum temporale or temporal plane of the superior temporal gyrus | | .08 | .05–.11 | < .001 |
| Mus. Peaceful | 62, -8, 4 | R Lateral aspect of the superior temporal gyrus | | .14 | .10–.18 | < .001 |
| Mus. Speed | 56, 2, -2 | R Planum polare of the superior temporal gyrus | | .26 | .19–.33 | < .001 |
| Mus. Speed | 62, -32, 13 | R Planum temporale or temporal plane of the superior temporal gyrus | | .24 | .16–.32 | < .001 |
| Mus. Speed | -62, 4, 1 | L Lateral aspect of the superior temporal gyrus | | .21 | .12–.30 | < .001 |
| Mus. Speed | -58, -38, 16 | L Planum temporale or temporal plane of the superior temporal gyrus | | .18 | .12–.24 | < .001 |
| Mus. Jitter | 58, -4, 7 | R Opercular part of the inferior frontal gyrus | | .17 | .14–.21 | < .001 |
| Mus. Jitter | -62, -10, 1 | L Lateral aspect of the superior temporal gyrus | | .13 | .08–.18 | .002 |
| Mus. Up/down | 56, -16, 10 | R Anterior transverse temporal gyrus (of Heschl) | | .17 | .13–.21 | < .001 |
| Mus. Up/down | -52, -22, 10 | L Anterior transverse temporal gyrus (of Heschl) | | .16 | .12–.20 | < .001 |
